# Cryo-EM reveals how ASX-173 inhibits human asparagine synthetase to activate the integrated stress response

**DOI:** 10.1101/2025.10.16.682859

**Authors:** Kirk A. Staschke, Nicholas T. Walda, Lucciano A. Pearce, Ronald C. Wek, Wen Zhu, Yuichiro Takagi

## Abstract

Targeting asparagine metabolism is a promising strategy for treating asparaginase-resistant acute lymphoblastic leukemia (ALL), sarcoma, and potentially other solid tumors. Here, we characterize the molecular mechanism by which a cell-penetrable small molecule, ASX-173, inhibits human asparagine synthetase (ASNS), the enzyme that catalyzes intracellular asparagine biosynthesis. ASX-173 reduces cellular asparagine levels, induces the integrated stress response (ISR), and reduces cell growth in HEK-293A cells. A cryo-EM structure reveals that ASX-173 engages a unique, hydrophobic pocket formed by AMP, Mg^2+^, and pyrophosphate in the C-terminal synthetase domain of ASNS, thereby enabling multivalent, high-affinity binding. Based on *in vitro* kinetic and thermal shift assays, we find that ASX-173 binds to the ASNS/Mg^2+^/ATP complex and is therefore a rare example of an uncompetitive enzyme inhibitor with potential therapeutic use. These findings provide a structural and mechanistic basis for targeting ASNS with small molecules, which have application in treating cancer and other human diseases.

## INTRODUCTION

Maintaining amino acid homeostasis is critically important for the overall health and viability of cells ^1-3^. The integrated stress response (ISR) pathway plays a crucial role in cell adaptation and survival to environmental stress ensuring sufficient amino acid uptake and synthesis to maintain homeostasis ^4^. A central feature of the ISR is stress-induced phosphorylation of the α subunit of eIF2 (p-eIF2α), triggering a reduction in global protein synthesis which conserves energy and nutrients and directs adaptive gene expression to restore amino acid homeostasis ^5,6^ (Fig.1a). The ISR is essential for many cancers, which inefficiently consume resources during uncontrolled cell division and tumorigenesis ^6^. Importantly, one critical ATF4 target gene encodes asparagine synthetase (ASNS), the enzyme that catalyzes the ATP-dependent biosynthesis of L-asparagine from L-aspartic acid using L-glutamine as a nitrogen source ^7^. The importance of asparagine metabolism for acute lymphoblastic leukemia (ALL) has been well-known for over 60 years ^8,9^, and asparaginase is a potent chemotherapeutic enzyme, which depletes this amino acid in leukemia cells that do not sufficiently express ASNS ^10,11^. Enhanced ASNS expression is correlated with cell proliferation in polycystic kidney disease (PKD) ^12^, sarcoma ^13^, a number of solid tumors ^14-20^, and the metastatic potential of breast cancer cells ^21^.

The critical role of ASNS for asparagine acquisition in cancers has driven efforts to identify potent, highly specific, small-molecule ASNS inhibitors ^22-26^. Prior studies have shown that analogs of either reaction intermediates or transition states (Supplementary Figs. 1-3) inhibit human ASNS with micromolar to nanomolar affinity ^23,26-28^. However, these ASNS inhibitors exhibit poor cytotoxicity, likely due to poor cell penetration. Recently, a cell-permeable ASNS inhibitor (ASX-173) was discovered ^29,30^ (Fig. 1b), which reduces the growth of a number of different cancer and leukemia cell lines. Moreover, ASX-173 potentiates the anti-cancer activity of L-asparaginase ^30^, opening up new possibilities for improved treatment of asparaginase-resistant ALL ^31^.

**Fig. 1.**
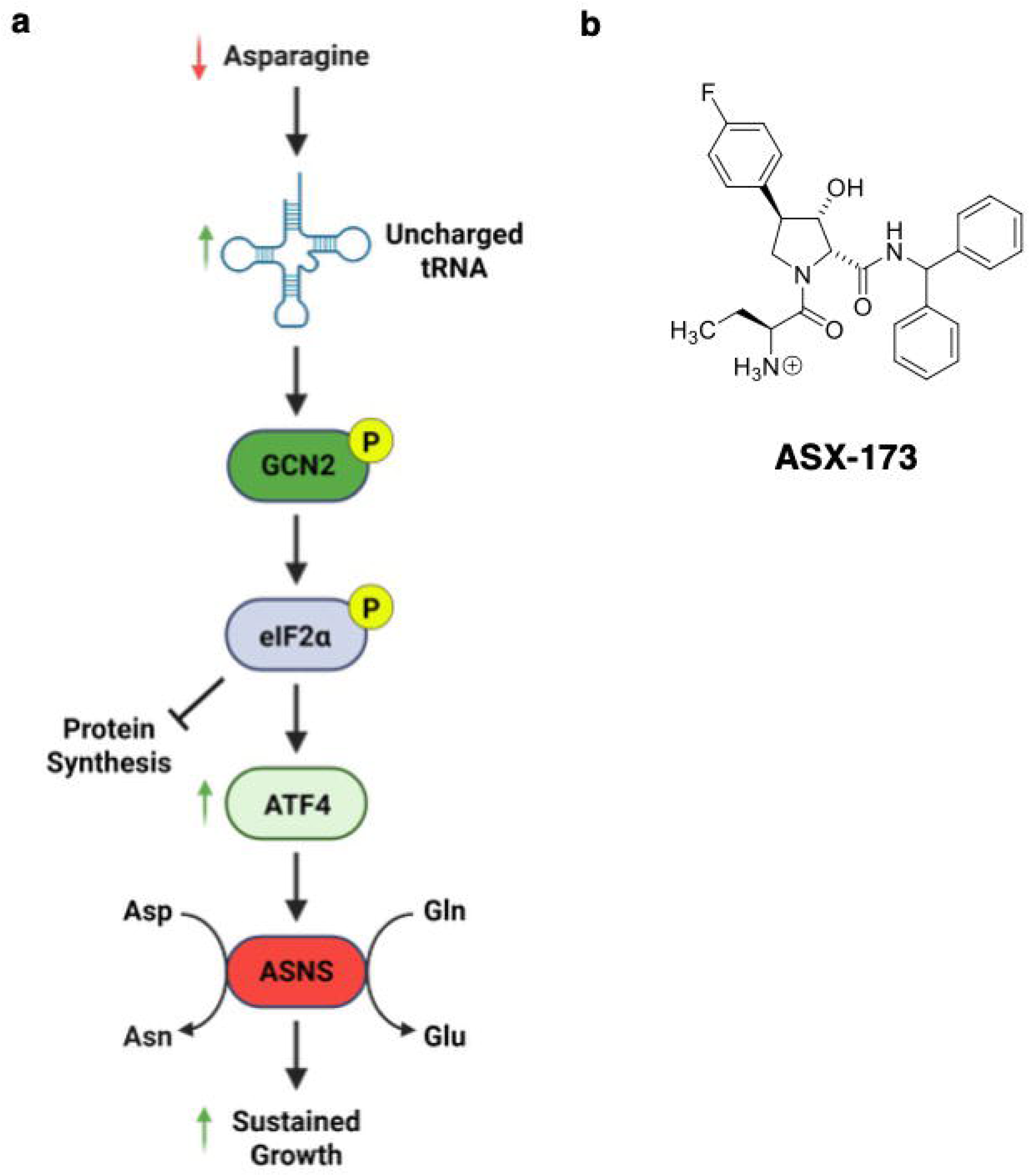
Schematic illustration of the ISR pathway and structure of ASX-173. **a** Asparagine depletion leads to increased phosphorylation eIF2α, resulting in attenuation of general protein synthesis and selective translation of ATF4. ATF4 controls ASNS expression, thereby restoring amino acid homeostasis. **b** The structure of ASX-173 is shown.

In this study, we use cellular, biochemical, and biophysical approaches to determine the molecular mechanism by which ASX-173 inhibits human ASNS. We show that ASX-173 inhibition of ASNS depletes cells of L-asparagine, and induces GCN2 eIF2 kinase and the ISR. Most importantly, we found that ASX-173 is a rare example of an uncompetitive enzyme inhibitor. This premise is supported by cryo-EM structure of the ASNS/ASX-173/Mg^2+^/AMP/pyrophosphate (PPi) complex at 2.58 Å resolution. Taken together, our findings provide a structural and mechanistic basis for the development of small-molecule drugs for treating PKD ^12^, sarcoma ^13^, and asparaginase-resistant ALL ^31^.

## RESULTS

### ASX-173 induces the ISR

Recently, a series of substituted hydroxy-pyrrolidine carboxamide compounds, including ASX-173 (Fig. 1b), were described that exhibit potent biochemical activity against ASNS and cytotoxicity against multiple cancer cell lines ^29,30^. We reasoned that targeted inhibition of ASNS in cells would create nutrient stress and induce the ISR. In support of this model, treatment of HEK-293A cells with ASX-173 resulted in substantial reduction in intracellular L-asparagine levels with a concomitant increase in other amino acids, presumably due to a reduction in global protein synthesis (Supplementary Fig. 4). Furthermore, ASX-173 treatment of the HEK-293A cells activated GCN2 as evidenced by increased p-GCN2-T899, phosphorylation of eIF2α, and expression of ATF4 in a concentration- and time-dependent manner (Fig. 2a and Supplementary Fig. 5a). This increase was comparable to a well-documented nutrient depletion mimic halofuginone (HF), an inhibitor of prolyl-tRNA charging and potent activator of GCN2 and the ISR ^32,33^. Consistent with activation of the ISR, treatment with ASX-173 also increased the expression of ISR-dependent genes, including *CHOP*, *GADD34*, and *TRIB3* (Fig. 2a, Supplementary Figs. 5a and 5b). It is noteworthy that ASNS, which is expressed at high basal levels in this cell line, is not substantially induced by ASX-173 treatment (Fig. 2a, Supplementary Figs. 5a and 5b). ASX-173 potently induced ATF4 transcriptional activity, and was reversed by the addition of physiological levels of L-asparagine to the culture media (Fig. 2b), indicating that selective inhibition of ASNS by ASX-173 induces the ISR.

**Fig. 2.**
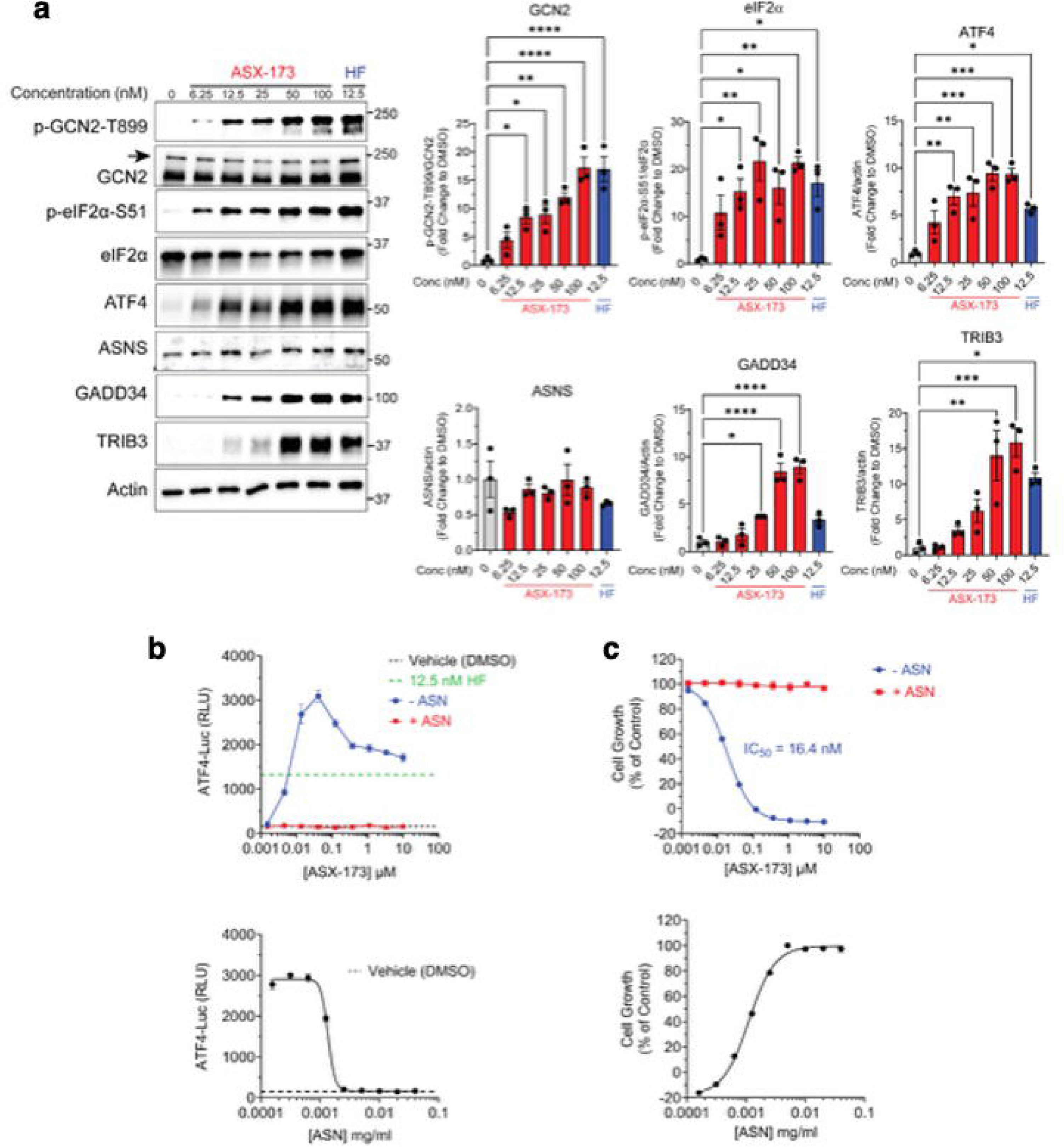
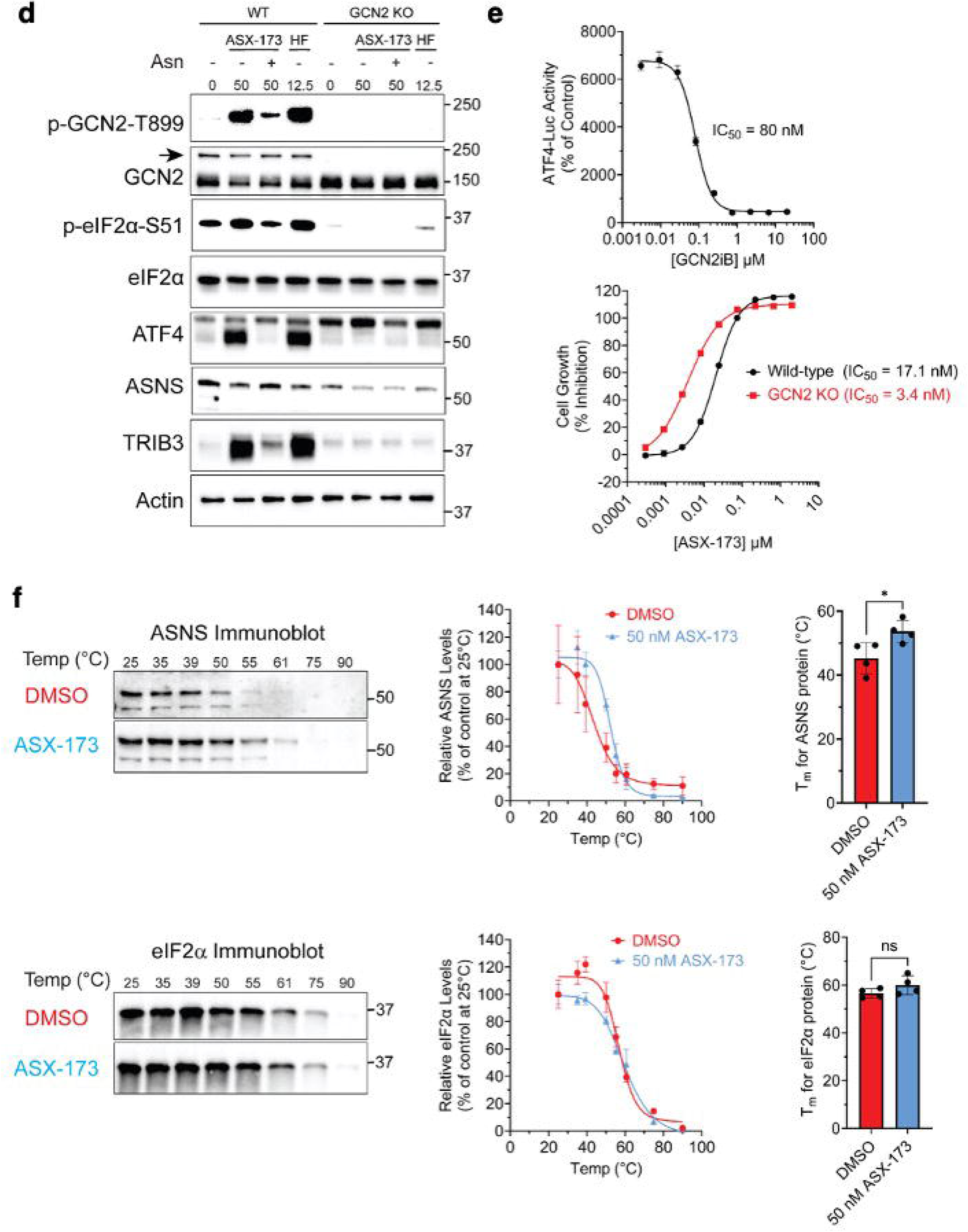
ASX-173 induces the ISR. **a** Lysates were prepared from 293A cells treated with ASX-173 (6.25 to 100 nM), HF (12.5 nM), or vehicle control (DMSO) for 6 hours, and immunoblot analysis was carried out using antibodies that recognize p-GCN2-T899, total GCN2, p-eIF2α-S51, total eIF2α, ATF4, ASNS, GADD34, TRIB3, or actin. An arrow marks the GCN2-specific band. Molecular weight markers are indicated in kilodaltons. Levels of the indicated proteins normalized to controls are shown in the bar graphs to the right. Statistical significance was determined using a one-way ANOVA with Tukey’s multiple comparisons (*n* = 3 biological replicates). **b** In the top panel, 293A reporter cells were treated with increasing amounts of ASX-173 (0.0015 µM to 10 µM), HF (12.5 nM), or vehicle (DMSO) as indicated for 6 hours (*n* = 3 biological replicates), and ATF4 transcriptional activity as measured by luciferase activity was determined. In the bottom panel, 293A reporter cells were treated with 50 nM ASX-173 in the presence of increasing levels of L-asparagine (0.0002 mg/ml to 0.04 mg/ml) or left untreated (DMSO control) as indicated for 6 hours (*n* = 3 biological replicates), and luciferase activity was determined to assess ATF4 transcriptional activity. **c** In the top panel, 293A reporter cells were treated with ASX-173 (0.0015 µM to 10 µM) or vehicle (DMSO) in the presence (+ ASN) or absence (- ASN) of L-asparagine (0.04 mg/ml) for 3 days (*n* = 3 biological replicates), and cell growth was determined using CellTiter-Glo^®^ 2.0 and is shown as a percentage of the untreated vehicle (DMSO) control. In the bottom panel, 293A reporter cells were treated with 50 nM ASX-173 in the presence of L-asparagine (0.0002 mg/ml to 0.04 mg/ml) or left untreated (DMSO control) as indicated for 3 days (*n* = 3 biological replicates), and cell growth was determined as described in the top panel above. **d** Lysates were prepared from 293A WT or 293A GCN2 KO cells treated with ASX-173 (50 nM) in the absence or presence of L-asparagine (Asn), HF (12.5 nM), or vehicle control (DMSO) for 6 hours, and immunoblot analysis was carried out using antibodies as described above in panel **a**. An arrow marks the GCN2-specific band. Molecular weight markers are indicated in kilodaltons. **e** In the top panel, 293A cells were treated with 50 nM ASX-173 in the presence of GCN2iB (0.003 µM to 20 µM) or vehicle (DMSO). Luciferase activity was determined after 6 hours of treatment and is indicated as a percentage of the DMSO control-treated samples. In the bottom panel, 293A WT or 293A GCN2 KO cells were treated with increasing concentrations of ASX-173 (0.0003 µM to 2.0 µM) or vehicle (DMSO) for 3 days (*n* = 3 biological replicates). Cell growth was determined as described in panel **c** and is shown as a percentage of vehicle (DMSO) control. **f** 293A cells were treated with 50 nM ASX-173 or vehicle (DMSO) for 1 hour (*n* = 4 biological replicates). Protein lysates were prepared and subjected to a thermal gradient as indicated, and soluble protein was analyzed by immunoblot for ASNS and eIF2α. Representative images of the immunoblots for ASNS and eIF2α are shown on the left. In the middle panel, the amount of soluble ASNS or eIF2α is plotted as a percentage of the levels present at 25 °C, and the melting temperature (T_m_) was calculated as described in the Materials and Methods section. In the right panel, the calculated T_m_ is plotted for each treatment group, and statistical significance was determined using a Welch’s unpaired two-tailed t-test. For all panels, there are error bars that indicate the standard error of the mean (SEM), and statistical significance is indicated as: ns; not significant; *, *P* ≤ 0.05; **, *P* ≤ 0.001; ***, *P* ≤ 0.001; and ****, *P* ≤ 0.0001.

Treatment of HEK-293A cells with ASX-173 significantly reduced cell growth, which was rescued by the addition of exogenous L-asparagine (Fig. 2c). Activation of GCN2 and induction of the ISR by ASX-173 were also reversed by the addition of L-asparagine and were completely blocked by deletion of the *GCN2* gene (Fig. 2d), indicating that activation of the ISR by selective inhibition of ASNS is GCN2-dependent. Furthermore, deletion of the *GCN2* gene substantially reduced ASNS expression, supporting the idea that *ASNS* is a key target gene of the ISR. Consistent with these findings, increased ATF4 transcriptional activity was inhibited by GCN2iB ^34^, a selective inhibitor of GCN2 (Fig. 2e, top panel). Interestingly, cells lacking GCN2 were more sensitive to ASNS inhibition by ASX-173. (Fig. 2e, bottom panel). Taken together, our results indicate that selective inhibition of ASNS by ASX-173 reduces intracellular L-asparagine levels selectively and potently induces GCN2-dependent activation of the ISR to maintain amino acid homeostasis.

We next tested whether ASX-173 directly engages ASNS in cells using a thermal protein profiling (TPP) assay ^35-37^. HEK-293A cells were treated with 50 nM ASX-173 or vehicle (DMSO) for 1 hour. Cell lysates were prepared and subjected to a temperature gradient from 25 °C to 90 °C, and the soluble protein levels were determined by immunoblotting for either ASNS or eIF2α as a control (Fig. 2f, left panel). Plots of the soluble protein levels of ASNS and eIF2α following thermal profiling were used to calculate the melting temperature (T_m_) between treatment groups (Fig. 2f, middle panel). Treatment with ASX-173 increased the T_m_ of ASNS from 45 °C to 54 °C, with no change in the thermal stability for eIF2α, indicating that ASX-173 binds to ASNS in cells (Fig. 2f, right panel). These results support the idea that ASX-173 directly binds to ASNS to inhibit its enzymatic activity.

### ASX-173 is an uncompetitive inhibitor with respect to Mg^2+^/ATP

Prior studies of bacterial glutamine-dependent asparagine synthetases (AS-B) have shown the presence of multiple intermediate states of enzyme-substrate, enzyme-intermediate, and enzyme-product complexes during catalytic turnover (Supplementary Fig. 6) ^38-40^. These states, including the apoenzyme, are all potential targets for ASX-173, with the situation being compounded further by the fact that ASNS is a bidomain enzyme, with each domain containing its own active site (Supplementary Fig. 7) ^41,42^. Thus, ammonia released from L-glutamine in the N-terminal glutaminase domain is translocated via an intramolecular tunnel to the C-terminal synthetase active site, where it undergoes reaction with a β-aspartyl-AMP intermediate formed from L-aspartate and Mg^2+^/ATP ^43^. In consequence, ASX-173 might impair catalysis at the N-terminal glutaminase active site, at the C-terminal synthetase active site, or by disrupting the interdomain tunnel that facilitates ammonia transfer.

To address how ASX-173 inhibits ASNS, we expressed and purified recombinant human ASNS following our published protocol (see Methods and Supplementary Fig. 8) ^41,42,44^ and determined IC_50_ values for ASX-173 under a variety of conditions. Rather than employing ASNS-mediated inorganic pyrophosphate (PPi) production as an indirect measure of asparagine formation, we directly determined the rate of L-asparagine formation and L-glutamate production using an HPLC-based end-point assay (Supplementary Fig. 9) ^38^. This approach was used because Mg^2+^PPi can be formed by hydrolysis of ATP or the β-aspartyl-AMP intermediate, even if ASX-173 binds to the glutaminase site of the enzyme. Dose-response curves were generated by titrating ASX-173 into reaction mixtures containing L-glutamine, L-aspartate, Mg^2+^ATP, and ASNS, with the percent inhibition (relative to control reactions containing 0.1% DMSO in place of the inhibitor) being evaluated as a function of ASX-173 concentration (Fig. 3a).

**Fig. 3.**
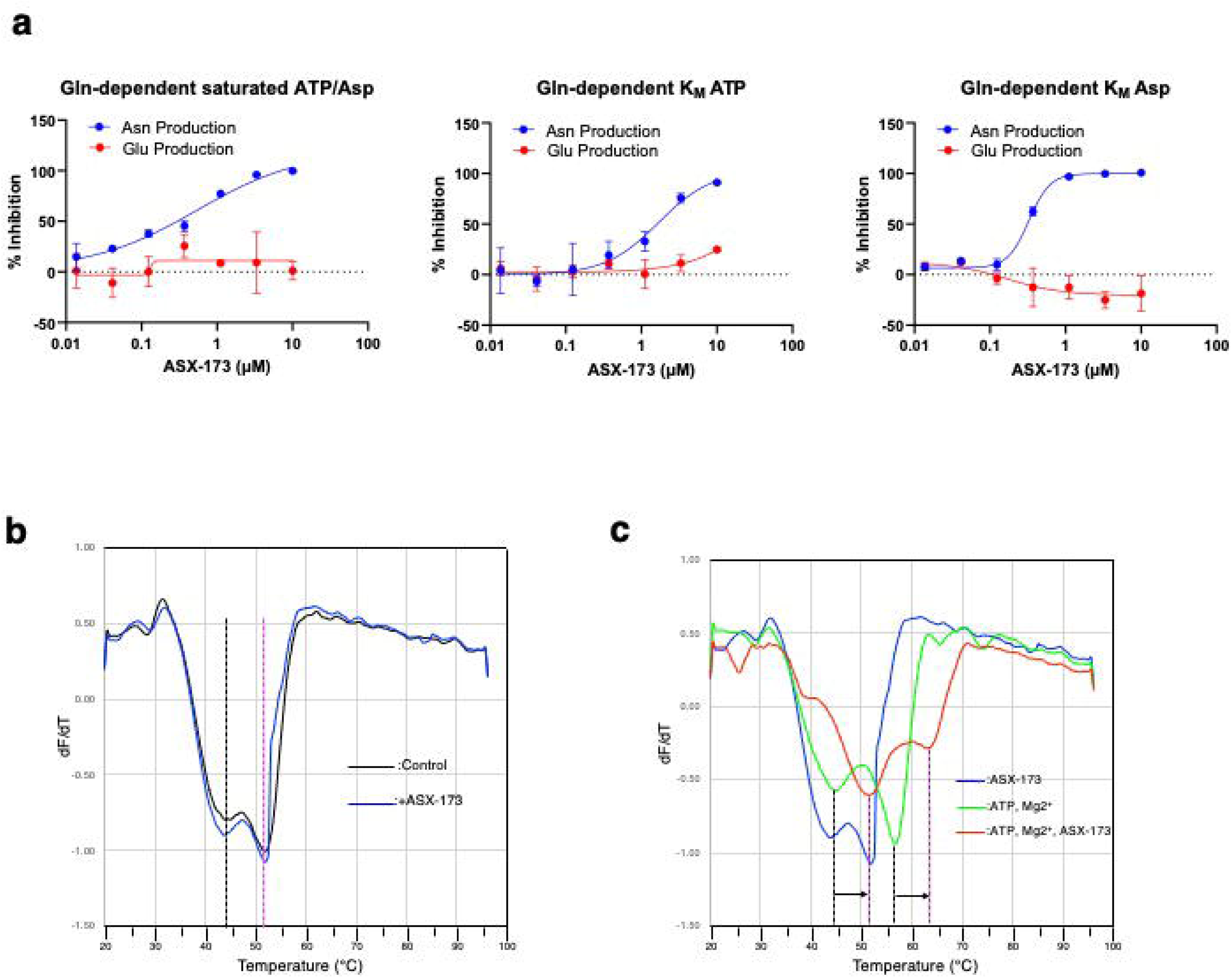
Effect of ASX-173 on kinetics for asparagine synthesis and the thermal stability of ASNS. **a** Dose-response inhibition of human ASNS by ASX-173. The production of L-asparagine (Asn) and L-glutamate (Glu) was quantified using an HPLC-based endpoint assay under L-glutamine (Gln)-saturated conditions with varying concentrations of ASX-173. The percentage inhibition of ASNS activity was determined under three substrate conditions: (left) saturated ATP and Asp, (middle) ATP at K_M_ and saturated Asp, and (right) Asp at K_M_ and saturated ATP. Percent inhibition was calculated relative to control reactions containing 0.1% DMSO instead of the inhibitor. Data represent mean ± SD from duplicated experiments. **b** The impact of ASX-173 on the thermal stability of recombinant ASNS was evaluated by differential scanning fluorimetry (DSF). ASNS (6 µg per assay) was incubated at room temperature for 1 hour and subjected to DSF. Control (buffer only, black), and ASX-173 at ten-fold molar excess (blue). Two dotted lines indicate two Tm: T_m1_ (black) and T_m2_ (magenta). **c** ASNS was incubated with 5 mM ATP with 10 mM Mg^2+^ (green), or with 5 mM ATP with 10 mM Mg^2+^ plus ASX-173 at ten-fold molar excess (green) prior to DSF. Dotted lines indicate Tm values and arrows indicate T_m_ shift.

When all three substrates are present at saturating concentrations, ASX-173 inhibited L-asparagine production with an IC_50_ value of 0.5 ± 0.2 μM (Fig. 3a), and glutamate production was unaffected. This result suggests that ASX-173 directly impairs asparagine production rather than perturbs *in situ* ammonia production, supporting the idea that the ASX-173 binding site is located in the C-terminal domain of ASNS. We therefore examined the effects of lowering the Mg^2+^ATP concentration to its apparent K_M_ value while maintaining L-glutamine and L-aspartate at saturating concentrations. Under these conditions, the IC_50_ value for the inhibition of L-asparagine production by ASX-173 increased more than three-fold even though the Hill slope remained relatively unchanged (Fig. 3a). This behavior is consistent with ASX-173 being an uncompetitive inhibitor with respect to ATP, meaning that it binds to a form of the enzyme produced after Mg^2+^ATP binds to the synthetase active site and not to the free enzyme as assumed in a previously reported kinetic analysis ^29^. A modest decrease in the rate of L-glutamate formation at higher concentrations of ASX-173 was seen under these conditions. Given that glutaminase activity is not inhibited by ASX-173 when all substrates are present at saturating concentrations, it is unlikely that the inhibitor directly targets the N-terminal glutaminase active site.

It is noteworthy that when both L-glutamine and Mg^2+^ATP were at saturating concentrations, the IC_50_ value of ASX-173 was unchanged by lowering L-aspartate from a saturating concentration to that corresponding to the apparent K_M_ value (Fig. 3a). Prior work on AS-B showed that L-aspartate binds to the enzyme/Mg^2+^ATP complex ^38^. Assuming that human ASNS employs a similar kinetic mechanism to AS-B, then competitive inhibition would be observed if L-aspartate and ASX-173 bind to the ASNS/Mg^2+^ATP complex in a mutually exclusive manner. On the other hand, if ASX-173 binds to either the ASNS/aspartate/Mg^2+^ATP or ASNS/β-aspartyl-AMP/Mg^2+^PPi complex, then inhibition should be uncompetitive with respect to L-aspartate. The unchanged IC_50_ value described here might therefore result from ASX-173 binding to a catalytic intermediate formed later in the kinetic mechanism, leading to L-asparagine ^38^. Our experiments also showed that glutamate production is slightly increased at higher inhibitor concentrations, in a similar manner to that observed for an *N*-methylsulfoxamine-based, transition-state analog of ASNS (Supplementary Fig. 1) ^26^. In that study, it was concluded that inhibitor binding at the active site stimulates glutaminase function via long-distance conformational changes. Based on our kinetic data, we conclude that ASX-173 interferes with L-asparagine production in the C-terminal active site by engaging one or more enzymatic states formed after the Mg^2+^ATP binding.

To test our interpretations of the *in vitro* kinetic results further, and as a necessary prelude to structural studies, we used a thermal shift assay ^45,46^ to assess the binding of ASX-173 for human ASNS under a variety of conditions. This assay not only measures how the melting temperature (T_m_) of the enzyme is affected by small-molecule ligands but also provides information on the number of folded forms that are present in solution. Thus, two T_m_ values, 44 °C (T_m1_) and 52 °C (T_m2_), were observed for the purified human ASNS when thermal denaturation was carried out for apo enzyme (Fig. 3b). Although other explanations are possible, we suggest that the first transition (T_m1_) arises from dimer dissociation at the N-terminal domain^42^. The second transition (T_m2_) therefore likely represents unfolding of the human ASNS monomer. Unexpectedly, the addition of a ten-fold excess of ASX-173 to the free enzyme (3.5 µM ASNS *vs*. 35 µM ASX-173) did not alter the melting profile (Fig. 3b). Indeed, the first derivative curves (dF/dT) for ASNS in the presence and absence of ASX-173 were virtually identical, suggesting that the inhibitor does not bind to the apo-enzyme. This finding is consistent with the results of our kinetic assays (Fig. 3a). Thus, if ASX-173 bound to the apo enzyme, we would expect the inhibitor to be competitive with respect to Mg^2+^ATP because the ASNS/Mg^2+^ATP complex is the first species formed in the ordered series of intermediate complexes needed for the production of L-asparagine (Supplementary Fig. 6) ^38^. We note that these thermal shift assay results raise concerns about the reported estimates of inhibitor binding, which are based on an incorrect kinetic model ^29^.

Given that ASX-173 does not bind to free human ASNS, we hypothesized that ASX-173 binding must depend on the presence of one, or more, of the substrates or products. As ATP and Mg²⁺ are essential for catalytic activity, we evaluated their effect on ASX-173 binding. Thermal shift assay showed that mixing human ASNS with ATP and MgCl_2_ increased the T_m2_ from 52 °C to 57 °C while leaving T_m1_ unchanged at 44 °C (Fig. 3c). More importantly, treating the enzyme with 35 µM ASX-173 in the presence of 5 mM ATP and 10 mM MgCl_2_ shifted both transitions upward so that T_m1_ and T_m2_ values of 52 °C and 63 °C were observed, respectively (Fig. 3c). These increased melting temperatures confirm that ASX-173 binds to either the ASNS/Mg^2+^ATP complex, or to an intermediate containing the products formed by Mg^2+^ATP hydrolysis. To resolve this issue, we performed a titration experiment in which the molar ratio of ASX-173 to ASNS was varied (1.25, 2.5, 5.0, and 10) (Supplementary Fig. 10). An upward T_m_ shift induced by ASX-173 in the presence of Mg^2+^ATP was observed even at the lowest molar ratio of 1.25, consistent with the observed sub-micromolar IC_50_ determined by steady-state kinetics (Fig. 3a). Finally, we repeated these experiments using AMP and Mg^2+^PPi, the products of Mg^2+^ATP hydrolysis, in place of Mg^2+^ATP, alongside a positive control, to determine whether these reaction products yielded a similar T_m_ shift profile. Compared with the positive control (Supplementary Fig. 11a), these results obtained with AMP and Mg^2+^PPi differed notably. No change was observed in T_m1_, while T_m2_ showed a modest increase from 52 °C to 54 °C in the presence of AMP and Mg^2+^PPi alone, and from 54 °C to 55 °C when ASX-173 was included with AMP and Mg^2+^PPi (Supplementary Fig. 11b). These results indicate that AMP and Mg^2+^PPi cannot fully substitute for Mg^2+^ATP in stabilizing ASX-173 binding to ASNS, suggesting that ATP hydrolysis could be essential for ASX-173 binding.

### ASX-173 binds to the synthetase site of human ASNS

Given these biochemical observations, we next employed single-particle cryo-EM to determine how ASX-173 binds within the ASNS/Mg^2+^ATP complex. Recombinant human ASNS was incubated with ASX-173 at a 1:2 molar ratio for 1 hour at room temperature in the presence of Mg^2+^ and ATP before cryo-EM grid preparation (see Methods). An initial dataset was collected on a 200 kV Glacios microscope and yielded a map at 3.14 Å resolution. Model building was guided by our previously determined cryo-EM structure of wild-type human ASNS (PDB: 8SUE)^42^ (Supplementary Figs. 12–14, Table 1). Additional densities not accounted for by the apo-ASNS structure were observed within the synthetase active site, corresponding to ASX-173 and suggesting the presence of AMP and Mg^2+^PPi (Supplementary Fig. 15). Despite the overall quality of the map being sufficient to position AMP, PPi, Mg^2+^, and ASX-173, however, some ambiguities remained.

To resolve these uncertainties, we collected additional data in super-resolution mode on a 300 kV Titan Krios microscope. Subsequent image processing (Methods) produced an improved map at 2.58 Å resolution (Figs. 4a and 4b; Supplementary Figs. 16–18, Table 1). In this higher-resolution map, all ligands were clearly visualized at the C-terminal synthetase site of each monomer (Fig. 4), thereby allowing unambiguous placement of ASX-173, AMP, and PPi into the extra densities, along with two Mg^2+^ ions (Fig. 4d) situated within the active site (Fig. 4). The presence of AMP, PPi, and Mg^2+^ was not entirely surprising in light of our thermal-shift data (Figs. 3b, 3c). Their presence in the structure indicates that ASX-173 binding may promote ATP hydrolysis by the enzyme.

**Fig. 4.**
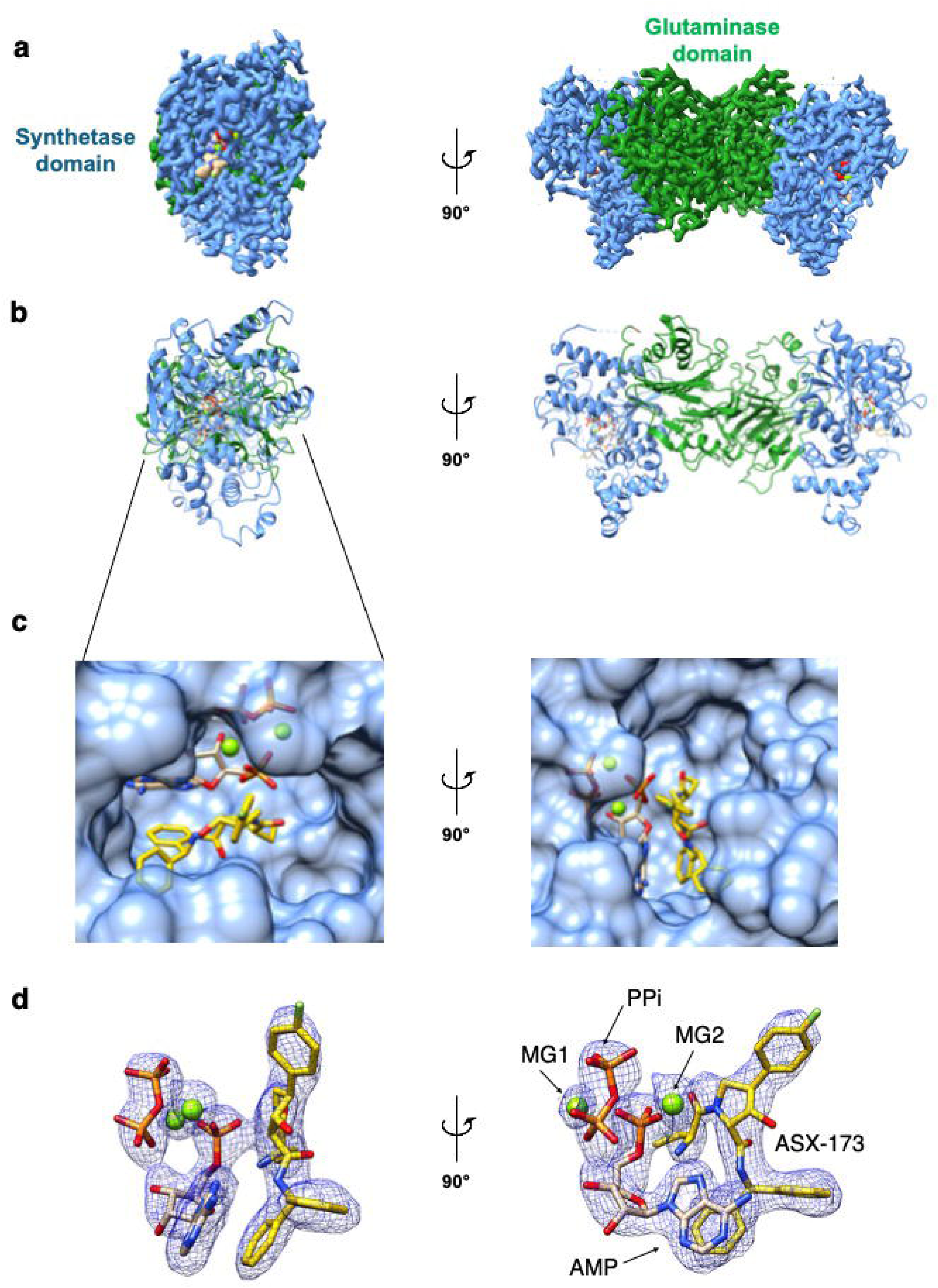
Cryo-EM structure of human ASNS in complex with ASX-173, AMPMg^2+^, PPi. **a** Cryo-EM density map of human ASNS/ASX-173/AMP/Mg^2+^PPi complex determined at 2.58 Å resolution. The glutaminase domain is colored in forest green and the C-terminal synthetase domain is colored in cornflower blue. Side view of ASNS dimer is on the left and the front view is on the right. The extra densities are seen in the synthetase site. **b** ribbon representation of human ASNS (coloring is identical to that used in **a**). **c** Close-up view of the synthetase site represented as surface highlighting the bound ligands: ASX-173, AMP, PPi, and Mg^2+^ ions. **d** Structure of how ligands fit together into the extra density in two different views. ASX-173 is colored in gold, two Mg^2+^ ions are colored in green. MG1, MG2 are indicated.

### Overall interactions of ASX-173, AMP, PPi, and Mg**^2+^** with human ASNS

Our cryo-EM structure shows that ASX-173 binds in close proximity to AMP and PPi within a composite binding pocket formed by ASNS. The inhibitor is anchored primarily by a network of hydrogen bonds to AMP and by extensive hydrophobic and π-interactions with residues of ASNS. No direct contacts are detected between ASX-173 and PPi. The detailed features of these intermolecular interactions, which collectively define a stable and well-organized ASNS/ASX-173/AMP/Mg^2+^PPi complex, are described below.

### AMP, PPi, Mg**^2+^**, and ASNS interactions

As predicted from bioinformatic analyses, Mg^2+^PPi interacts with residues in the conserved ‘PP-loop’ motif comprising Ser-257, Gly-258, Gly-259, Leu-260, and Asp-261 (the canonical SGGxD motif) (Fig. 5a) ^47^. The spatial arrangement of PPi, the PP-loop, and AMP closely resembles that observed in other enzymes containing evolutionarily related ATP pyrophosphatase domains, which include GMP synthetase ^48^ and argininosuccinate synthetase ^49,50^ (Supplementary Fig. 19). AMP is stabilized through hydrogen bonding involving the phosphate groups of PPi, as well as residues Ile-287 and Ser-362 (Fig. 5a). Notably, our EM map revealed an additional density near the active site that has not been reported in any previous X-ray or cryo-EM structures of ASNS ^41,42^. We assigned this density to a second Mg^2+^ ion, thereby raising the intriguing possibility that catalysis may involve a two-metal mechanism. The two Mg^2+^ ions (designated MG1 and MG2) (Fig. 4d) are coordinated by the carboxylate side chains of Asp-261 and Asp-367, together with the phosphate groups of AMP and PPi (Fig. 5a). This metal-bridging network appears critical for neutralizing the local negative charge and stabilizing the interactions among ASNS, AMP, and PPi.

**Fig. 5.**
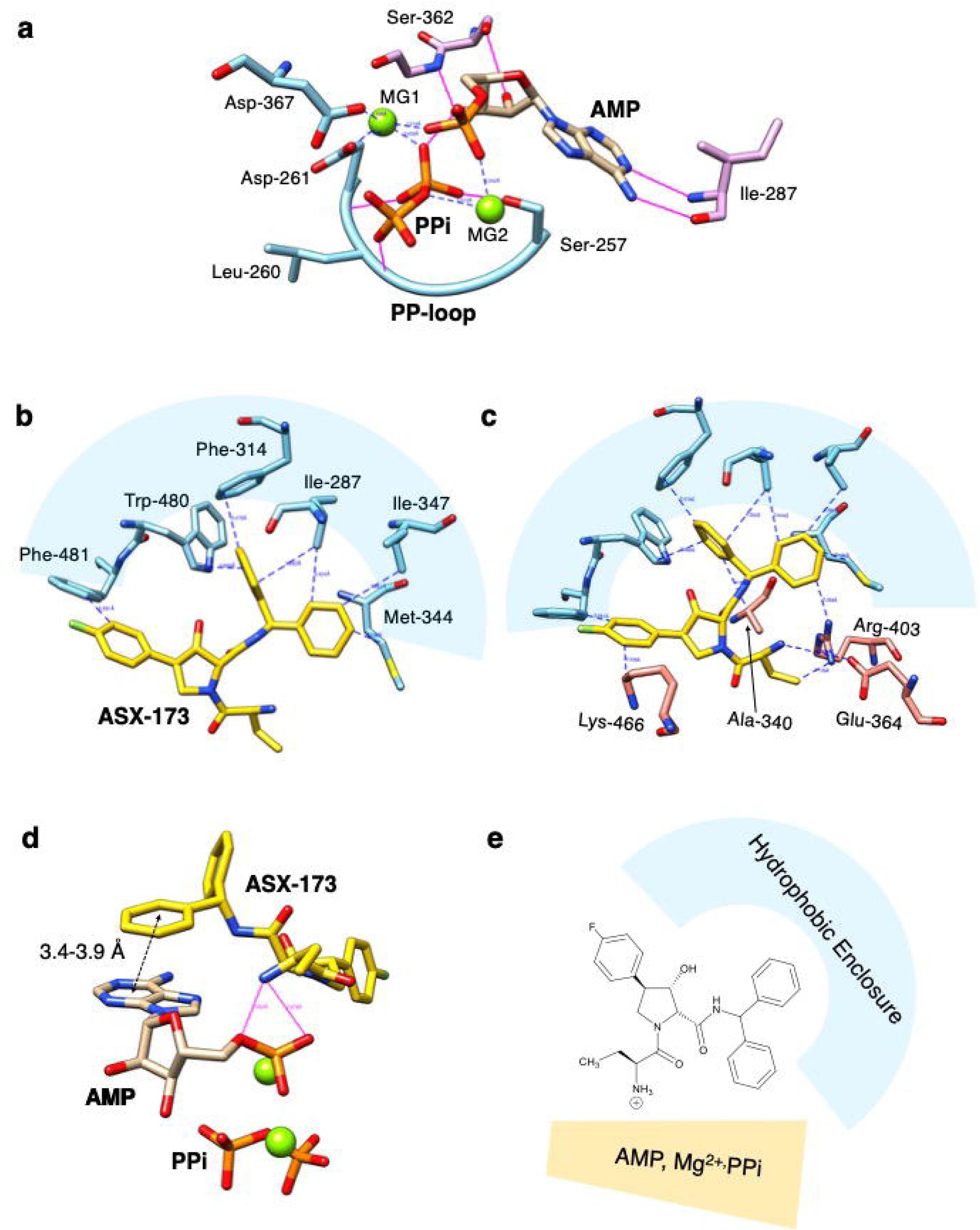
Structural basis of ligand coordination and ASX-173 binding at the synthetase site. **a** Coordination network of AMP, PPi, Mg^2+^, and the conserved PP-loop motif (SGGLD) (sky blue) of ASNS together with additional residues Ile-287 and Ser-362 (pink) that further stabilize AMP. Two Mg²⁺ ions (MG1 and MG2) are coordinated by Asp-261, Asp-367, and the phosphate groups of AMP and PPi, forming a metal-bridging network that neutralizes negative charge and stabilizes the active-site complex. **b** Hydrophobic enclosure formed by Ile-287, Phe-314, Met-344, Ile-347, Trp-480, and Phe-481 that binds the aromatic and aliphatic regions of ASX-173 (colored in gold) through van der Waals and π-π, and π-alkyl interactions. **c** Polar and cation-π interactions contributed by Arg-403 and Lys-466, together with Ala-340 and Glu-364, further position ASX-173. **d** Phosphate-directed hydrogen bonds and π-π stacking interactions between AMP and ASX-173. The phosphate group of AMP hydrogen-bonds with the amine atoms of ASX-173 (2.5–2.9 Å), while the adenine base engages in π-π and π-alkyl contacts (3.4-3.9 Å) with the aromatic moiety of ASX-173. Mg²⁺ ions are colored in green. **e** Illustration of the ASNS/ASX-173 AMP/Mg^2+^/PPi interactions. A hydrophobic enclosure and the coordination network of AMP, PPi, and Mg^2+^ stabilizes ASX-173 binding to ASNS. This tripartite arrangement underlies the uncompetitive inhibition mechanism, wherein ASX-173 preferentially binds the enzyme in its AMP-bound, post-hydrolysis state.

### ASX-173, AMP, PPi, and ASNS interactions

The structural model reveals that ASX-173 is bound within a composite pocket formed jointly by residues from the synthetase domain of ASNS and the bound AMP molecule. The inhibitor is positioned by a combination of hydrophobic, π-mediated, and phosphate-directed polar interactions that collectively anchor it at the active site. On one side of the pocket, residues Ile-287, Phe-314, Met-344, Ile-347, Trp-480, and Phe-481 form a hydrophobic enclosure that accommodates the aromatic and aliphatic portions of ASX-173 through van der Waals and π-π and π-alkyl interactions (Fig. 5b). Arg-403 and Lys-466 contribute additional cation-π contacts; Ala-340 and Glu-364 form polar interactions, albeit at distances of 3.6-4.3 Å, shaping the contours of the binding cavity without engaging in direct hydrogen bonding (Fig. 5c).

On the opposite face, AMP reinforces ASX-173 binding through a network of phosphate-directed hydrogen bonds and π-π stacking with the adenine base. The phosphate group of AMP forms hydrogen bonds with the amine of ASX-173 (2.5-2.9 Å), while the adenine ring interacts with the aromatic moiety of ASX-173 via π-π stacking ^51^ (3.4–3.9 Å) (Fig. 5d). These interactions orient the inhibitor within the synthetase domain in the presence of Mg^2+^PPi, effectively bridging the active site residues with ATP-derived intermediates. No direct contacts were observed between ASX-173 and PPi within 4 Å. Together, the hydrophobic enclosure contributed by ASNS and the phosphate-anchored hydrogen-bonding network provided by AMP generate a composite, amphipathic binding pocket that stabilizes ASX-173 through synergistic nonpolar, π-π stacking, and polar interactions. This tripartite arrangement, comprising ASNS residues, AMP, and ASX-173, likely underlies the uncompetitive inhibitory mechanism of ASX-173, in which the inhibitor preferentially associates with the enzyme in its AMP-bound, post-hydrolysis state.

Even in the presence of all ligands, interpretable density is missing for two loop regions (residues 201-220 and 465-475) and the C-terminal tail (residues 542-560). These segments are also absent in all previous X-ray and cryo-EM structures of human ASNS and the R142I variant ^41,42^, suggesting that these regions are intrinsically disordered.

Finally, as indicated by our previous findings ^42^, Arg-142 plays a key regulatory role in gating ammonia transfer between the N-terminal glutaminase domain and the C-terminal synthetase site through conformational changes in its side chain. 3D variability analysis (3DVA) of the ASX-173–bound form of ASNS revealed that Arg-142 adopts a conformation that likely blocks ammonia diffusion from the N-terminal to the C-terminal site (Supplementary Fig. 20). These observations suggest that the inhibitory effect of ASX-173 might extend beyond the synthetase active site to include disruption of the ammonia channel connecting the two catalytic domains.

### ASX-173 does not bind to evolutionarily related ATP pyrophosphatases

ASNS shares key active site features with other ATP pyrophosphatases, such as GMP synthetase (GMPS) and argininosuccinate synthetase (ASS1), including a conserved spatial arrangement of PPi, the PP-loop, and AMP (Supplementary Fig. 19) ^48,50^. Because ASX-173 binds to ASNS through phosphate-directed hydrogen bonds and π-π stacking with AMP, we tested whether GMPS or ASS1 could also serve as targets for this small molecule. Recombinant human GMPS and ASS1 were expressed in insect cells, purified (Supplementary Fig. 8) and analyzed using a thermal shift assay ^45,46^ under four conditions: (i) apo enzyme, (ii) +ASX-173 alone, (iii) +Mg^2+^ATP alone, and (iv) +Mg^2+^ATP/ASX-173. In the case of GMPS, the T_m_ remained unchanged at 52 °C across all conditions (Supplementary Fig. 21a). For ASS1, Mg^2+^ATP raised the T_m_ from 47 °C to 52 °C, but ASX-173 (either alone or with Mg^2+^ATP) had no effect (Supplementary Fig. 21b). These results show that ASX-173 does not bind to GMPS or ASS1, suggesting that it binds only to human ASNS even in the presence of other cellular pyrophosphatases. This hypothesis is supported by the fact that exogenous L-asparagine can reverse the effects of ASX-173 on cell growth and proliferation (Fig. 2).

### The cryo-EM model is consistent with observed structure-activity relationships

ASX-173 is only one of a series of analogs (Supplementary Fig. 22) that have been characterized for both their ability to inhibit ammonia-dependent asparagine production by human ASNS and cytotoxicity to a number of cancer cell lines ^29^. Despite our concerns about the kinetic model used in the patent ^29^, plotting the apparent inhibition constants against cytotoxicity towards a Jurkat T cell line still yields a reasonable correlation between activity and affinity (Supplementary Fig. 23). For example, when ASX-173 and a diastereoisomer (in which the reversed absolute stereochemistry of the fluorophenyl and hydroxyl substituents) are mixed in a 1:1 ratio, the cytotoxicity remains essentially unchanged, despite the inhibition constant being increased two-fold. This finding is consistent with only one of the two diastereoisomers having the correct stereochemistry to position the hydroxyl group correctly relative to the Asn-478 side chain, an interpretation that is reinforced by the ability of analogs **4** and **23** to inhibit the enzyme. Interestingly, methylating the hydroxyl substituent (compound **22**) only reduces affinity by an order of magnitude, presumably because the side chain amide moiety can reorient to maintain the hydrogen bonding interaction. Our model also reveals an electrostatic interaction between the protonated amine in ASX-173 and the phosphate of AMP. The importance of this salt bridge is demonstrated by the poor ability of **46**, in which the amine is converted into an amide, to inhibit ASNS and kill cells. On the other hand, the importance of the fluorophenyl substituent for binding and cytotoxicity is not clearly defined by this set of analogs.

## DISCUSSION

Our results demonstrate that ASX-173 is an uncompetitive inhibitor of human asparagine synthetase (ASNS), binding specifically to the ASNS/Mg^2+^ATP complex rather than the apo enzyme. This finding is unexpected and mechanistically distinct from previously reported ASNS inhibitors, which mimic the β-aspartyl-AMP transition state and bind to the unliganded enzyme ^23,24,26^. In kinetic terms, uncompetitive inhibition reduces the pool of catalytically competent enzyme-substrate complexes, thereby reducing catalysis and effectively lowering intracellular asparagine levels. The resulting amino acid deficiency activates the ISR, consistent with our cellular data and the established role of the GCN2-ATF4 pathway in sensing nutrient limitation. Thus, this kinetic model provides a direct mechanistic link between ASNS inhibition, reduced asparagine biosynthesis, and ISR activation.

ASX-173 was discovered by cell-based screening, rather than the use of enzyme-based assay screening against the apo enzyme. This cell-based method likely enabled the identification of a rare substrate-dependent uncompetitive inhibitor, analogous to isofagomine, which binds to the glucosyltransferase domains of *Clostridium difficile* toxin only in the presence of UDP-glucose ^52^. It has also been reported that purine nucleoside phosphorylase inhibitor binding may be dependent on the presence of bound phosphate ^53^. These observations emphasize the value of physiological screening strategies in uncovering inhibitors that exploit enzyme-substrate complexes.

The therapeutic significance of the biochemical and structural work reported here is underscored by recent work from Lorenzi *et al.*, showing that ASX-173 potentiates the activity of L-asparaginase and significantly delays leukemia xenograft growth in mice ^30^. Together, our work and these studies validate ASNS as a tractable anti-cancer target whose inhibition disrupts both asparagine metabolism and stress-response signaling. Structurally, our cryo-EM model of the ASNS/ASX-173/AMP/Mg^2+^PPi complex reveals that ASX-173 interacts with ASNS primarily through hydrophobic, cation/π-π stacking with ASNS and AMP, and minimal direct hydrogen bonds. Our thermal shift and kinetic assays suggest that direct inhibitor engagement is promoted by the presence of catalytic products, e.g., AMP and Mg^2+^PPi, and is primarily driven by the substrate, Mg^2+^ATP. Our study establishes a mechanistic and structural framework for designing next-generation ASNS inhibitors and highlights uncompetitive, substrate-dependent inhibition as a promising strategy for targeting asparagine metabolism as a metabolic vulnerability in cancers.

## METHODS

### Reagents

HEK-293A-ATF4-luc and GCN2 knock-out reporter cells ^54^ were cultured in DMEM (Corning, Cat. #A4899101) supplemented with 10% fetal bovine serum (HyClone, Cat. #SH30071) and 1 µg/mL puromycin (MP Biomedicals, Cat. #194539). Cell lines were maintained at 37°C in a fully humidified atmosphere containing 5% CO_2_, used at early passage for all experiments, and tested to be free of mycoplasma contamination. ASX-173 or compound 17 ^29,30^. was synthesized at Sun-shine Chemical (Wuhan, China), and stock solutions (10 mM) were prepared in DMSO. The structure and purity of ASX-173 were confirmed by ^1^H-NMR and LC-MS (Supplementary Figs. 2 and 3), respectively, by the vendor and in the Chemical Genomics Core at the Indiana University School of Medicine. Halofuginone (Cat. #13370) was purchased from Caymen Chemical Company (Ann Arbor, MI), GCN2iB (Cat. HY-112654) was purchased from MedChemExpress (Monmouth Junction, NJ), and L-asparagine (Cat. A0884-100G) was purchased from Sigma-Aldrich (St. Louis, MO).

### Cell culture and treatments

For growth assays, HEK-293A reporter cells were seeded into 96-well plates at 2,500 cells per well and treated as indicated for 3 days. Cell growth was monitored using CellTiter-Glo 2.0 Reagent (Promega) and was calculated relative to a day 0 control. For ATF4 transcriptional activity assays, 293A reporter cells were seeded into 96-well plates at 15,000 cells per well and treated as indicated for 6 hours. Luciferase activity was measured using Bio-Glo™ luciferase assay reagent (Promega).

### Immunoblot analysis

293A reporter cells were seeded at 300,000 cells per well in 6-well or 35 mm plates in normal culture medium and treated as indicated. Whole cell extracts from cultured 293A reporter cells were prepared in 1% sodium dodecyl sulfate (SDS) lysis buffer supplemented with 1× Halt Protease and Phosphatase Inhibitor Cocktail (Thermo Scientific, Cat. #1861281). Lysates were boiled for 10 minutes, sonicated, and subjected to centrifugation at 20,000 × g to remove insoluble material. Protein concentrations were determined by DC Protein Assay (Bio-Rad, Cat. #5000112) using bovine serum albumin (BSA) as the protein standard. Equal amounts of protein were separated on SDS-polyacrylamide gel electrophoresis gels, transferred to nitrocellulose membranes, and used for immunoblot analysis. The primary antibodies used were as follows: phospho-GCN2-T899 (Abcam Cat. #ab75836, RRID:AB_1310260), total GCN2 (Cell Signaling Technology Cat. #3302, RRID:AB_2277617), phospho-eIF2α-S51 (Abcam Cat. #ab32157, RRID:AB_732117), total eIF2α (Cell Signaling Technology Cat. #5324, RRID:AB_10692650), ATF4 (Cell Signaling Technology Cat. #11815, RRID:AB_2616025), or custom rabbit polyclonal antibody which was prepared against full-length recombinant human ATF4 protein and affinity purified, ASNS (Cell Signaling Technology, Cat. #20843S), TRIB3 (Abcam Cat. #ab75846, RRID:AB_1310768), GADD34 (Proteintech Cat. #10449-1-AP), and β-actin (Sigma-Aldrich Cat. #A5441, RRID:AB_476744). Immunoblot signals were visualized by enhanced chemiluminescence (ECL) using Clarity Western ECL Substrate (Bio-Rad Laboratories, Inc., Cat. #170-5060) or SuperSignal West Femto Maximum Sensitivity Substrate (Thermo Fisher Scientific, Cat. #34094), and images were captured using a ChemiDoc Imaging System (Bio-Rad Laboratories, Inc., RRID:SCR_019037). Pre-stained protein standards (Bio-Rad Laboratories, Inc., Cat. #1610377) were included on each immunoblot, captured as a combined image file, and are indicated in each immunoblot panel.

### qRT-PCR analysis

293A reporter cells were seeded into 100 mm plates at 2×10^6^ cells per plate and treated as indicated. Following treatment, total RNA was isolated using TRIzol LS Reagent (Invitrogen, Cat. #10296010) and quantified using a Nanodrop Spectrophotometer (Thermo Fisher Scientific). Total RNA (2 µg) was converted to cDNA using a High-Capacity cDNA Reverse Transcription Kit (Thermo Fisher Scientific, Cat. #4368813) according to the manufacturer’s protocol. RNA samples were diluted 10-fold in nuclease-free water (Cytiva, Cat. #SH30538.02), and PCR amplification was carried out using PowerUp SYBR Green Master Mix (Thermo Fisher Scientific, Cat. #A25724) using an Applied Biosystems QuantStudio5 PCR System. The primers used for measuring ATF4, ASNS, CHOP, GADD34, TRIB3, and ACTB (actin) are listed elsewhere (Supplementary Table 2). The relative abundance of each transcript was calculated using the ΔΔCT method with ACTB (Actin) serving as an internal control, and the data are presented normalized to each vehicle control (DMSO) group.

### Target engagement assay

Engagement of ASNS by ASX-173 in cells was determined using a cell-based thermal shift assay [47-49]. 293A reporter cells (2 × 10^6^) were cultured in 100 mm dishes in standard growth medium and treated with vehicle (DMSO) or 50 nM ASX-173 for 1 hour. Following treatment, the cells were rinsed with cold PBS and harvested in 350 µl of lysis buffer (40 mM HEPES, pH 7.5, 200 mM NaCl, 0.4% NP-40) containing HALT™ Protease and Phosphatase Inhibitor Cocktail (Pierce, Cat. #1861281). Cell lysates were prepared by 30 cycles of sonication in a 4 °C water bath for 30 seconds, followed by centrifugation at 300 x *g* for 5 minutes. The protein concentration of the resulting supernatant was determined using a DC Protein Assay (Bio-Rad, Cat. #5000112) using bovine serum albumin (BSA) as the protein standard. The resulting lysates were adjusted to 1 mg/ml using complete lysis buffer, and 50 µl aliquots were subjected to a heat gradient consisting of 25 °C for 2 minutes, 25.0 °, 35.0 °, 39.3 °, 50.1 °, 55.2 °, 60.7 °, 74.9 °C, or 90.0 ° for 3 minutes, followed by 2 minutes at 25 °C, and cooling to 4 °C. Following the temperature gradient, lysates were centrifuged at 20,000 × g for 30 minutes at 4 °C to remove insoluble material. Equal volumes of the remaining supernatant were mixed with Laemmli buffer, heated to 95 °C, and analysed by immunoblotting for ASNS and Actin, as described above. The specific protein bands were quantified using Image Lab Software (Bio-Rad Laboratories, Inc.), and the melting temperature (T_m_) was determined from the inflection point of 4-parameter fitted curves using GraphPad Prism.

### Amino acid measurements

Amino acid levels were measured in cultured cell lysates by liquid chromatography–tandem mass spectrometry (LC-MS) as previously described ^55,56^. Briefly, 293A reporter cells (500,000) were cultured in normal growth medium and treated with vehicle (DMSO) or ASX-173 (50 nM) in standard growth for 4 hours. The cells were rinsed briefly with cold phosphate-buffered saline (PBS) and then harvested by scraping in 0.1% formic acid in methanol vortexed briefly, and subjected to three cycles of freezing in liquid N_2_ for 2 min, followed by thawing at 42°C for 10 min. Samples were further homogenized using a handheld mortar and pestle and clarified by centrifugation at 10,000 × g at 4 °C for 10 minutes. The resulting lysates were passed through a Captiva Non-Drip Lipid spin column (Agilent Technologies) by centrifugation at 2,000 RPM at 4 °C for 10 minutes. Clarified lysates were analysed using LC–MS/MS with stable isotope dilution. Lysates were spiked with a mixture of amino acid isotopic internal standards and deproteinated using a solution of 90% acetonitrile,10 mM ammonium formate, and 0.15% formic acid. Clarified lysates were analysed using a biphasic LC-MS/MS approach, first with a Hydrophilic Interaction Liquid Chromatography (HILIC) separation ^57^, followed by a mixed-mode chromatographic separation ^58^. Quantification of amino acid levels was determined by fitting response ratios to an eight-point calibration curve generated using verified reference material for each of the 20 amino acids quantified.

### Expression and purification of recombinant ASNS

Expression and purification of human ASNS was performed as previously described ^41^. Briefly, a 1LL culture of Sf9 insect cells (1.5 × 10^6^Lcells/mL) was infected with baculovirus encoding either WT ASNS at a multiplicity of infection (eMOI) of 4.0 ^44^. Infected cells were incubated at 27 °C for 72 hours, harvested by centrifugation, flash-frozen in liquid nitrogen, and stored at - 80 °C. Cell pellets were lysed in 50 mM Tris-HCl, pHL8.0, containing 500 mM NaCl. The lysate was clarified by ultracentrifugation and purified by nickel-affinity chromatography. The His_x6_-tag was removed by TEV protease digestion, followed by further purification using a 1 mL HiTrap Q anion exchange column (Cytiva, Wilmington, DE). Peak fractions were pooled, dialyzed against the buffer containing 50 mM Tris-HCl, pHL8.0, 200 mM NaCl and 5 mM DTT, and concentrated to 8.4 mg/mL using a spin column (MilliporeSigma, St. Louis, MO). Protein concentration was determined by Bradford assay, and aliquots of purified ASNS were stored at -80 °C. Purified ASNS was subjected to 4-12% NuPAGE (Invitrogen) and stained with Coomassie blue, which indicates high quality of recombinant ASNS (Supplementary Fig. 9).

### Expression and purification of recombinant human GMP synthetase (GMPS) and argininosuccinate synthetase (ASS1)

The open reading frames (ORFs) of human GMPS (UniProtKB: P49915) and ASS1 (UniProtKB: P00966) were codon-optimized, synthesized, and subcloned into the EcoRV site of pUC57 (GenScript, Piscataway, NJ, USA) to generate pUC57-BamHI-GMPS-HindIII and pUC57-BamHI-ASS1-HindIII plasmids. The BamHI–HindIII fragments were subsequently inserted into the corresponding sites of the pKL-10His-3C vector ^59^, which encodes an N-terminal His10 tag followed by a 3C protease cleavage site ^60^, yielding pKL-10His-3C-GMPS (pYT1703) and pKL-10His-3C-ASS (pYT1704) transfer vectors.

Recombinant baculoviruses expressing GMPS or ASS1 were generated in Sf9 cells using these vectors as described previously ^44^. Protein expression was optimized using the TEQC method ^44^. In brief, 200 mL cultures of Sf9 cells (Expression Systems, Inc.) were infected at an estimated multiplicity of infection (eMOI) of 4 and incubated for 96 h at 27 °C. Cells were harvested, flash-frozen in liquid nitrogen, and stored at -80 °C until use. Cell pellets from 200 mL cultures were lysed in 50 mL lysis buffer (50 mM Tris-HCl, pH 8.0, 500 mM NaCl, 5 mM imidazole, 5% glycerol, 0.01% NP-40, and 5 mM β-mercaptoethanol) supplemented with protease inhibitors (6 mM leupeptin, 0.2 mM pepstatin A, 20 mM benzamidine, and 10 mM PMSF) ^44^. Lysates were stirred for 30 min at 4 °C and centrifuged at 100,000 × *g* for 30 min. The supernatant (6 mL for GMPS and 3 mL for ASS1) was applied to HIS-Select® Nickel Affinity Gel (Millipore-Sigma) pre- equilibrated with lysis buffer. After loading, the resin was washed sequentially with 5 column volumes (CV) of lysis buffer and 5 CV of lysis buffer containing 15 mM imidazole (without NP-40).

From this point, purification procedures diverged because high-salt conditions were necessary to stabilize ASS1, as reported previously ^61^. For GMPS, the resin was further washed with 10 CV of Buffer B+200 (50 mM Tris-HCl, pH 8.0, 200 mM NaCl, 15 mM imidazole, 5% glycerol, and 5 mM β-mercaptoethanol), and the protein was eluted with elution buffer (50 mM Tris-HCl, pH 8.0, 200 mM NaCl, 300 mM imidazole, 5% glycerol, and 5 mM β-mercaptoethanol). For ASS1, the resin was washed with 5 CV of the same buffer (15 mM imidazole, no NP-40) and eluted using 50 mM Tris-HCl pH 8.0, 500 mM NaCl, 300 mM imidazole, 5% glycerol, and 5 mM β-mercaptoethanol. Eluted fractions were pooled, and buffer exchanged using PD-10 columns (Cytiva) to Buffer B+200 for GMPS and lysis buffer without NP-40 for ASS1, followed by concentration with 10 kDa cutoff Amicon filters to 3.8 mg/mL (GMPS) and 2.0 mg/mL (ASS1), as determined by Bradford assay. Purified GMPS and ASS1 were subjected to 4-12% NuPAGE (Invitrogen) and stained with Coomassie blue, which indicated high quality of recombinant GMPS and ASS1 (Supplementary Fig. 9).

### Steady-state kinetic assays

An HPLC-based enzyme inhibition assay was developed to assess the dose-response of ASX-173 under varying substrate conditions. Serial dilutions of ASX-173, prepared in a final concentration of 0.1% DMSO, were added to reaction mixtures at 37 °C containing 100 mM EPPS buffer, pH 8.0, 2 mM DTT, 10 mM MgCl_2_, and varying concentrations of substrates, including ATP (saturated = 5 mM, K_M_ = 0.08 mM), L-aspartate (saturated = 12.5 mM, K_M_ = 0.38 mM), and L-glutamine (20 mM). Reactions were initiated by adding ASNS and were quenched with glacial acetic acid after 30 minutes. The pH was then adjusted to 9.0 using NaOH and Na_2_CO_3_ solution before derivatization with a saturated solution of 1-fluoro-2,4-dinitrobenzene (DNFB) in DMSO. The derivatization reaction was carried out at 50 °C for 45 minutes and subsequently quenched with glacial acetic acid. Samples were analysed by reverse-phase HPLC using a C18 column to separate DNFB-derivatized amino acids. Standard curves for asparagine and glutamate were generated by applying the same derivatization protocol to known concentrations of the amino acids. Peak areas were converted to concentrations using these standard curves, and percent inhibition was calculated from the fitted four-parameter curves using GraphPad Prism.

### Thermal shift assay

Thermal shift assays were performed to assess the thermal stability of human asparagine synthetase (ASNS), human GMP synthetase (GMPS) and human argininosuccinate synthetase (ASS1) in the presence or absence of ASX-173, ATP, and MgCl₂. Each 25 µL reaction was prepared in a 96-well qPCR plate using a buffer composed of 50 mM Tris-HCl, pH 8.0, 200 mM NaCl, 5% (v/v) glycerol, and 5 mM β-mercaptoethanol. Reactions contained 3.5 µM ASNS (6.2 µg), or GMPS (6 µg) or ASS1 (8 µg), 1× GloMelt™ dye (Biotium, Fremont CA), and the indicated combinations of ASX-173, 5 mM ATP, and 10 mM MgCl₂. Samples were incubated at room temperature for 1 hour prior to thermal scanning. In the experiment assessing the effect of AMP and PPi, 3.5 µM ASNS (6.2 µg), 5 mM AMP, 5 mM PPi, and 10 mM MgCl₂ were used.

Thermal unfolding was monitored on a Bio-Rad CFX Connect real-time PCR detection system. Fluorescence was recorded continuously using the SYBR Green channel as the temperature increased from 20L°C to 95L°C at a rate of 1.0L°C per minute. Data were acquired and analyzed using CFX Maestro software (Bio-Rad). All measurements were duplicated. Fluorescence singles were normalized to close to the maximum value in the same dataset. The T_m_ was determined from the peak of the first derivative of the fluorescence signal (dF/dT) plotted as a function of temperature.

### Cryo-EM sample preparation and data collection

A 10 mM stock solution of ASX173 in 100% DMSO was diluted with the buffer [50LmM Tris-HCl (pHL8.0), 200 mM NaCl, 5% glycerol, and 5 mM β-mercaptoethanol] to prepare a 1 mM working solution. Recombinant ASNS (8.4 mg/mL), ASX173 (1 mM), ATP (100 mM), and MgCl₂ (100 mM) were mixed with 10x Buffer [500 mM Tris-HCl (pHL8.0), 2M NaCl] and H_2_O at a final concentration of 48 μM ASNS, 100 μM ASX-173, 5 mM ATP, and 10 mM MgCl₂, corresponding to an approximate molar ratio of 2:1 (ASNS:ASX-173). The mixture was incubated at room temperature for 1 hour prior to cryo-EM grid preparation.

300 mesh UltrAuFoil R1.2/1.3 grids were glow-discharged for 1 minute with a current of 15mA in a PELCO easiGlow system before being mounted onto a Mark IV Vitrobot (FEI/Thermo Fisher Scientific). The sample chamber on the Vitrobot was kept at 4 °C with a relative humidity of 100%. 3.0 μL of the recombinant human ASNS/ASX173 complex at a concentration of 3.2 mg/ml was applied to the grid, which was then blotted from both sides for 4 seconds with blot force set at 0. After blotting, the grid was rapidly plunge-frozen into a liquid ethane bath cooled by liquid nitrogen.

The data was collected by a 200 kV Glacios transmission electron microscope (FEI/ThermoFisher Scientific) equipped with a Falcon4 camera operated in Electron-Event Representation (EER) mode ^62^. Movies were collected using EPU at 150,000x magnification (physical pixel size 0.93 Å) over a defocus range of -0.4 to −2.0 μm and a total accumulated dose of 40 e/ Å^2^. Camera gain reference was taken at the end of the run. A total of 4,680 movies were acquired. Each movie was split into 40 fractions during motion correction with EER upsampling factor = 1. After motion correction and CTF estimation, micrographs were manually curated based on relative ice thickness and CTF resolution fit such that 3,229 micrographs out of 4,680 were chosen and used for subsequent data processing.

High-resolution data was collected by a 300 kV Tithan Krios G4 transmission electron microscope (FEI/ThermoFisher Scientific) equipped with a K3 detector with BioQuantum Energy Filter (Gatan). Movies were collected at super-resolution mode using EPU at 105,000x magnification (physical pixel size 0.411 Å) over a defocus range of -0.8 to -1.8 μm and a total accumulated dose of 60 e/ Å^2^. During motion corrections, movies were binned 2×, resulting in dose-weighted micrographs in 0.822/pixel. After motion correction and CTF estimation, micrographs were manually curated based on relative ice thickness and CTF resolution fit such that 6,412 micrographs out of 8,184 were chosen and used for subsequent data processing.

### Cryo-EM data processing

The full cryo-EM data processing workflows are illustrated in (Supplementary Figs. 12 and 16). Motion correction, CTF-estimation, particle picking, 2D classification, *Ab-initio* 3D reconstruction, heterologous refinement, and non-uniform 3D refinement were performed in cryoSPARC v4.7.0^63^.

### 200 kV Glacios cryo-EM data processing and structure determination

Template for particle picking were generated from the apo-ASNS structure (PDB: 8SUE) using UCSF Chimera ^64^. As shown (Supplementary Fig. 12), template-based particle picking was performed, followed by three rounds of 2D classification to remove false positives and damaged particles. The cleaned particle set was then subjected to two rounds of *ab initio* reconstruction and heterogeneous refinement, yielding three distinct initial 3D maps. Particles contributing to the highest quality map among the three were selected for another cycle of *ab initio 3D* reconstruction and heterogeneous refinement, again producing three maps. The best of these maps, with an estimated resolution of 4.31 Å, was further refined using non-uniform refinement with global CTF, per-particle defocus, beam tilt corrections, and C2 symmetry, resulting in a 3.57 Å map. Particles from this map were grouped by beam shift and underwent global CTF refinement, followed by homogeneous refinement. The output map was further processed with another round of non-uniform refinement using global CTF, defocus, beam tilt corrections, and C2 symmetry, improving the resolution to 3.36 Å.

A total of 91,104 particles constituting the map were then re-extracted from the micrographs using a box size of 320 pixels. These re-extracted particles (88,344) were subjected to non-uniform 3D refinement with C1 symmetry. This was followed by a final round of non-uniform refinement with global CTF, defocus, beam tilt corrections, and C2 symmetry, yielding a final map at 3.14 Å resolution. Resolution was estimated based on Fourier Shell Correlation (FSC) using the gold-standard approach with a 0.143 cut-off criterion ^65-67^. The final density map was sharpened using DeepEMhancer ^68^. via the COSMIC^2^ platform ^69^. All EM maps were visualized using UCSF Chimera ^64^ and ChimeraX ^70^.

### 300 kV Titan Krios Cryo-EM data processing and structure determination

Template-based particle picking was performed using the same template generated from the apo-ASNS structure (PDB: 8SUE), as described above. As shown (Supplementary Fig. 16), initial particles were subjected to two rounds of 2D classification to eliminate bad particles. This particle set underwent four iterative rounds of *ab initio* reconstruction and heterogeneous refinement, yielding three initial 3D maps. Particles contributing to the two highest-quality maps were selected and subjected to a new round of *ab initio* reconstruction, followed by heterogeneous refinement and another round of *ab initio* reconstruction and heterogeneous refinement, generating three more 3D maps. From these, particles constituting the two best maps were further refined through *ab initio* reconstruction and heterogeneous refinement, yielding three 3D maps. The best of these maps, with a resolution of 3.37 Å, was refined using non-uniform refinement with C1 symmetry, resulting in a map at 2.91 Å resolution. This map was further refined using non-uniform refinement with global CTF correction, per-particle defocus and beam tilt corrections, and C2 symmetry, yielding a 2.79 Å map. Particles contributing to this map were re-extracted from the micrographs using a box size of 320 pixels. The re-extracted particles underwent an additional round of non-uniform refinement with C1 symmetry, followed by a final round of non-uniform refinement using global CTF, defocus, beam tilt corrections, and C2 symmetry. This yielded the final map at 2.58 Å resolution, estimated using Fourier Shell Correlation (FSC) with the gold-standard 0.143 cut-off criterion ^65-67^. The final density map was sharpened using DeepEMhancer ^68^ via the COSMIC^2^ platform ^69^. All cryo-EM maps were visualized using UCSF Chimera ^64^ and ChimeraX ^70^.

### Model building and refinement

The EM map was sharpened using the DeepEMhancer toolL^68^ via the COSMIC^2^ platform ^69^. and the resulting map was used for model building. Initial fitting was performed by placing the apo-ASNS crystal structure (PDB: 8SUE) into the sharpened EM map using rigid-body refinement in REFMACL^71^. This initial model was further refined through iterative rounds of real-space refinement in PhenixL^72^, followed by manual inspection and adjustment in Coot. Additional unmodeled densities, not accounted for by the apo-ASNS structure, were identified and attributed to three ligands: ASX-173, AMP, and PPi. Two additional densities near AMP and PPi were assigned as Mg^2+^ ions. Geometry restraints and coordinate files (CIF format) for ASX-173, AMP, and PPi were generated using the eLBOW module in Phenix ^73^. The complete model, including ligands and metal ions, was refined using real-space refinement in Phenix with the corresponding ligand restraints, followed by further manual adjustments in Coot. Final model refinement statistics are summarized in Supplementary Table 1.

### 3DVA of ASNS-ASX173-AMP-Mg**^2+^**PPi complex EM map

The EM map was subjected to 3D variability analysis (3DVA) in cryoSPARC v4.7.0 with 5 principal components filtered at 4 Å resolution with the mask used in a final non-uniform refinement ^74^. The movie frames were generated by the 3DVA display program in a simple mode in cryoSPARC v4.7.0. Out of 20 movie frames, frame 0 at one end, and frame 10 at the other end were subjected to variability refinement in Phenix ^75^. The model originated from frame 0 at one end and that from frame 19 at the other end were used for subsequent analysis.

### Data and Materials Availability

EM-derived structural data of the complex have been deposited in the PDB (PDB ID: 9YNS, 9YNT) and in the Electron Microscopy Data Bank (EMDB ID: EMD-73230, EMD-73232). Other data can be downloaded using links provided in the Supplementary Materials.

## Acknowledgments

This work was supported in part by grants from the NIH R35GM136331 (RCW) and R35GM160337 (WZ), the Department of Defense HT9425-24-1-0525 (RCW, KAS), Indiana University Simon Comprehensive Cancer Center P30CA082709 (KAS), the American Cancer Society IRG-22-147-37-06 (KAS), and the Indiana Clinical and Translational Sciences Institute (YT, KAS). The authors would like to acknowledge the Indiana University School of Medicine EM facility and NIH/NIGMS, S10 OD028723, for supporting this work. The authors thank Drs. Vago and Klose in the Purdue EM facility (RRID:SCR_025545) for assisting with data collections. Additional funds for the EM structure determinations were provided by the Indiana University School of Medicine (YT), and, in part, NIGMS award number R01GM111695 (YT) and the American Cancer Society award number DBG-23-1038947-01-IBCD (YT). The authors acknowledge Florida State University for additional financial support through start-up funds provided to W.Z.

## Author contributions

WZ, KAS and YT conceived of and designed the studies; NTW and KAS performed the cell-based studies; LAP and WZ performed enzyme purification, kinetic assay development, and data analysis; YT carried out enzyme expression, thermal shift measurements, and solved the cryo-EM structure of the ASNS/ASX-173/AMP/Mg^2+^PPi complex. WZ, KAS, and YT wrote the initial draft of the manuscript and supporting information, with subsequent review and editing by all other authors.

