## Supplementary Information for "Cryo-EM reveals how ASX-173 inhibits human asparagine synthetase to activate the integrated stress response"

**β-aspartyl-AMP analogs**

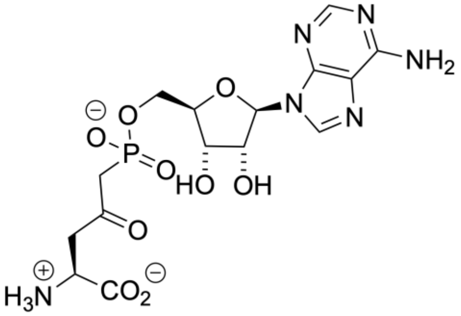

Biochim. Biophys. Acta **708**, 203-209 (1982)

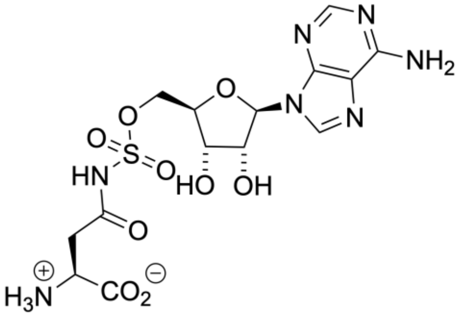

Org. Lett. **5**, 2033-2036 (2003)

**methylsulfoximine transition state analogs**

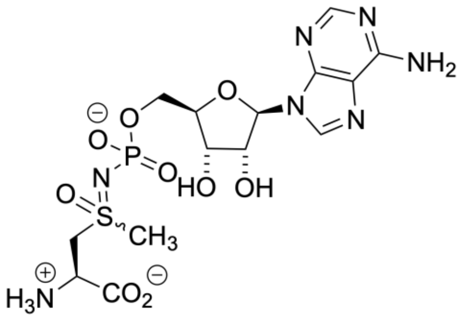

Chem. Biol. **13**, 1349-1347 (2006)

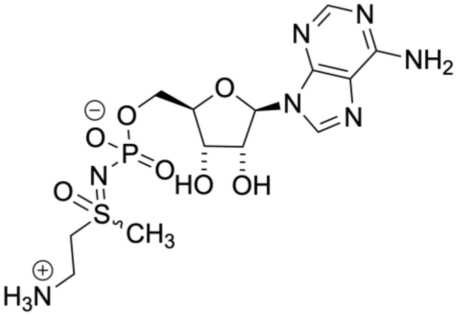

Bioorg. Med. Chem. **20**, 5915-5927 (2012)

**natural products**

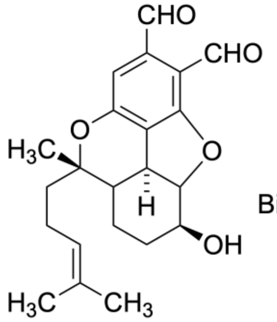

Bisabosqual A

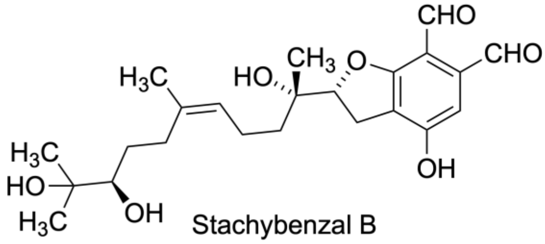

Stachybenzal B

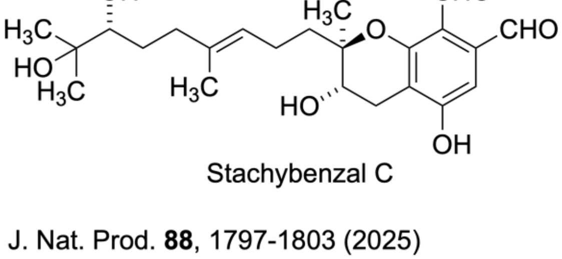

Stachybenzal C

Eur. J. Pharmacol. **960**, 176156 (2023)

J. Nat. Prod. **88**, 1797-1803 (2025)

**Supplementary Fig. 1. Known small molecule ASNS inhibitors.** First-generation inhibitors were unreactive analogs of β-aspartyl-AMP, an intermediate formed in the synthetase active site during catalytic turnover. More potent, second-generation inhibitors resembled the transition for the reaction of ammonia with β-aspartyl-AMP. Recent studies have identified a series of natural products that inhibit activity by covalent modification of a lysine residue in the C-terminal tail of the enzyme.

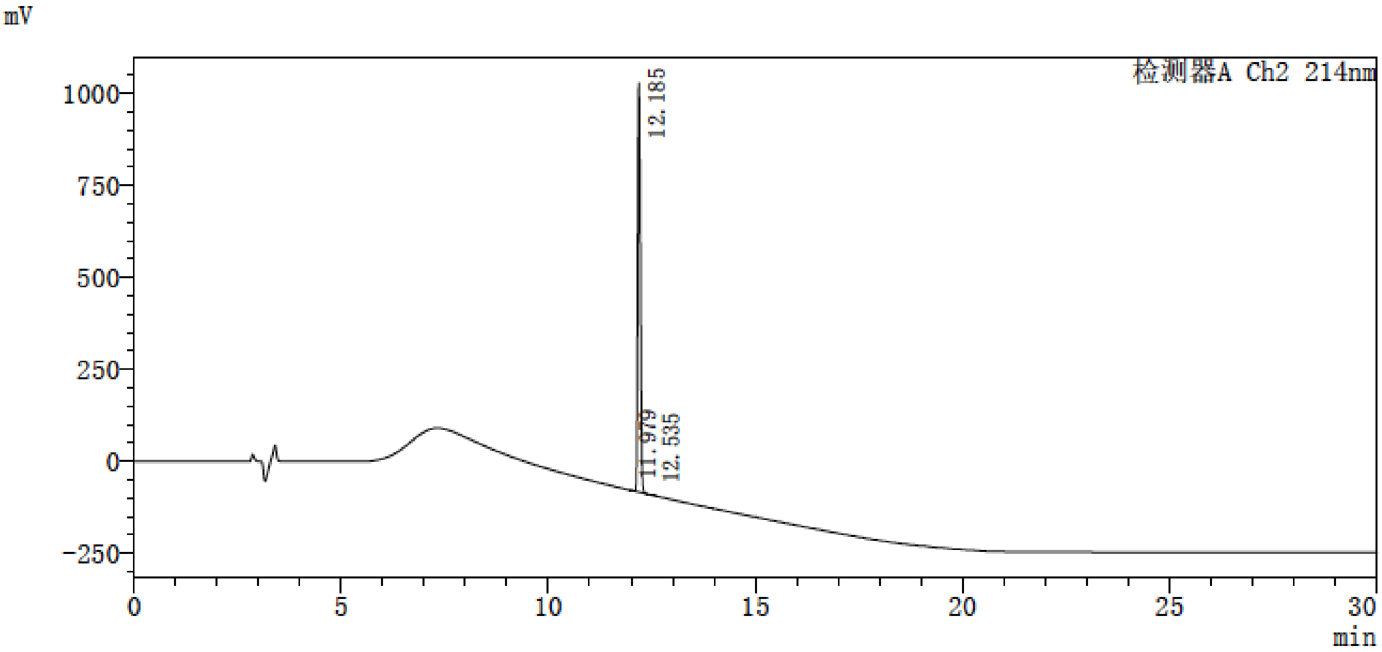


**Supplementary Fig. 2. HPLC chromatogram of ASX-173.** Chemical synthesis yields the ASNS inhibitor in > 98% purity.

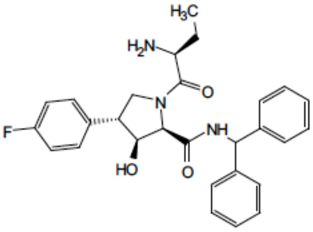


[1]H NMR (400 MHz, DMSO) δ 8.80 (dd, $J$ = 125.5, 8.3 Hz, 1H), 7.61 – 7.01 (m, 12H), 6.28 – 6.05 (m, 1H), 5.54 (dd, $J$ = 104.9, 5.9 Hz, 1H), 4.72 (dd, $J$ = 91.8, 7.8 Hz, 1H), 4.43 (dt, $J$ = 14.9, 7.0 Hz, 1H), 4.24 – 3.73 (m, 1H), 3.55 (q, $J$ = 9.2 Hz, 1H), 3.40 (d, $J$ = 10.5 Hz, 1H), 3.31 – 3.23 (m, 1H), 1.56 (d, $J$ = 20.8 Hz, 2H), 1.40 – 1.17 (m, 1H), 1.15 – 0.76 (m, 2H), 0.48 (t, $J$ = 7.3 Hz, 1H).

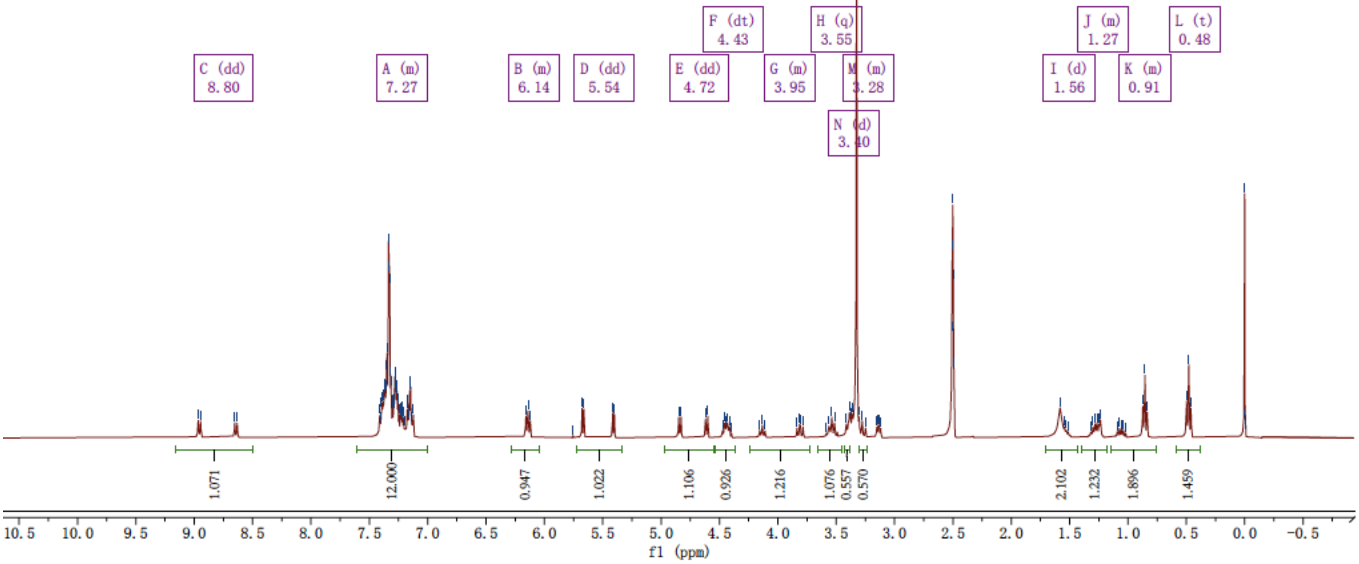


| | | | | | |
|---|---|---|---|---|---|
| **Sample Name** | | **Position** | P2-B1 | **Instrument Name** | Instrument 1 |
| **User Name** | | **Inj Vol** | 1 | **InjPosition** | |
| **Sample Type** | Sample | **IRM Calibration Status** | Success | **Data Filename** | 20241008 ASX-173.d |
| **ACQ Method** | Positive_AB_P2_0.6mLPerMin_7Min.m | **Comment** | | **Acquired Time** | 10/8/2024 2:14:12 PM (UTC-04:00) |

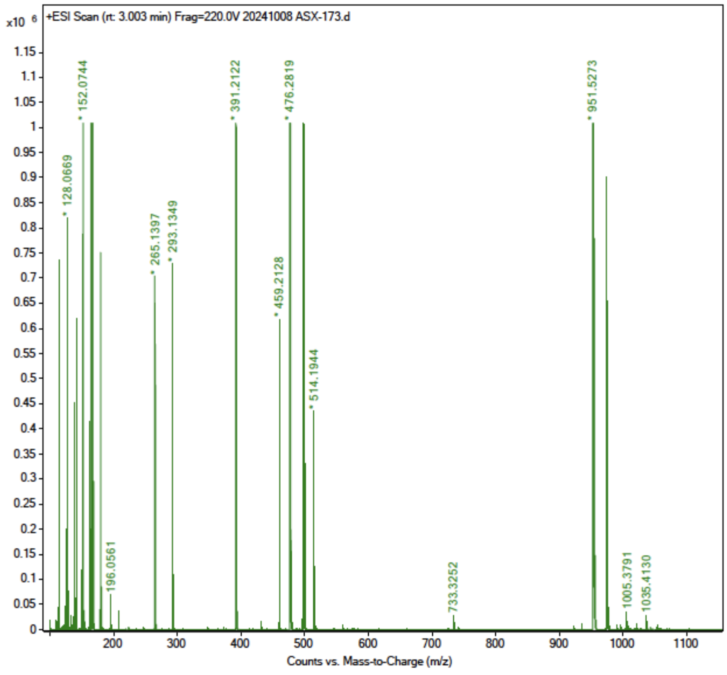


**Supplementary Fig. 3. Spectroscopic data for ASX-173.** (**top**) 400 MHz [1]H NMR spectrum of ASX-173 dissolved $d_6$-DMSO. (**bottom**) Electron ionization mass spectrum of ASX-173.

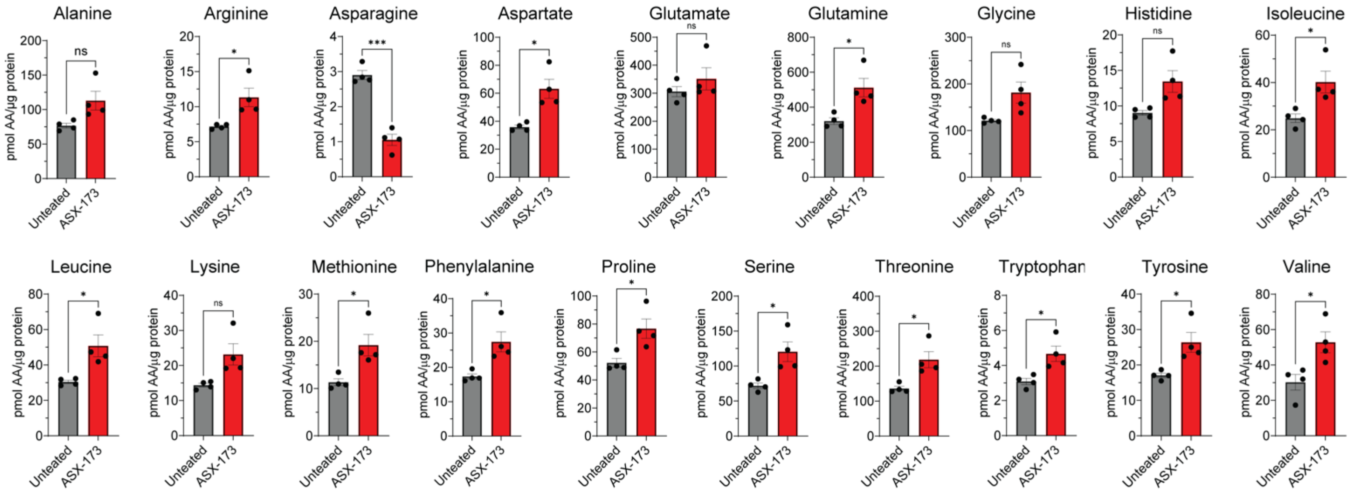


**Supplementary Fig. 4**. **ASX-173 reduces intracellular asparagine levels.** Levels of free amino acids were measured in 293A cells treated with 50 nM ASX-173 or vehicle (DMSO) for 6 hours (*n* = 4 biological replicates). Bar graphs indicate the amount of each amino acid in picomoles normalized to total protein. Note, cystine was not detected in this experiment. Error bars indicate the standard error of the mean (SEM). Statistical significance was determined using a Welch's unpaired two-tailed t-test and is indicated as: ns; not significant; *, $P \leq 0.05$; and ***, $P \leq 0.001$.

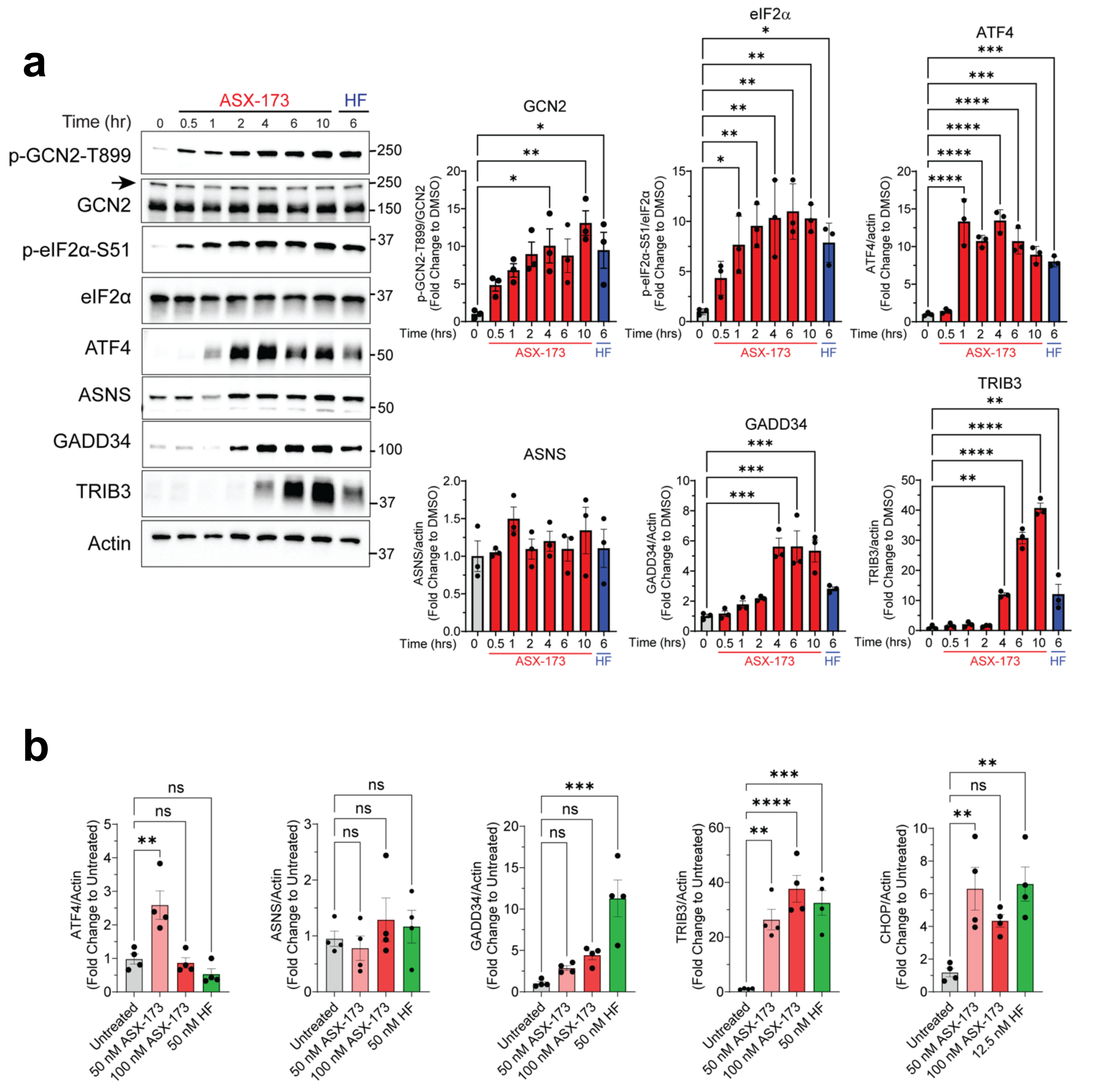


**Supplementary Fig. 5. ASX-173 induces the ISR and ISR-dependent gene expression. (a)** Lysates were prepared from 293A cells treated with 50 nM ASX-173 for 0.5 to 10 hours. Cells were treated with 12.5 nM HF, or left untreated (DMSO, 0 hour) as a control. Immunoblot analysis was carried out using antibodies that recognize p-GCN2-T899, total GCN2, p-eIF2α-S51, total eIF2α, ATF4, ASNS, TRIB3, or actin as described in Figure **2a**. An arrow marks the GCN2-specific band. Molecular weight markers are indicated in kilodaltons. Levels of the indicated proteins normalized to controls are shown in the bar graphs to the right. Statistical significance was determined using a one-way ANOVA with Tukey's multiple comparisons ($n$ = 3 biological replicates). **(b)** 293A cells were treated with 50 or 100 nM ASX-173, 50 nM HF, or vehicle (DMSO), and mRNA levels were measured for *ATF4*, *ASNS*, *GADD34*, *TRIB3*, *CHOP*, or *ACTB* by qRT-PCR, and relative levels are shown normalized to ACTB (actin). Statistical significance was determined using a one-way ANOVA with Tukey's multiple comparisons ($n$ = 4 biological replicates). For all panels, error bars indicate the standard error of the mean (SEM), and statistical significance is indicated as: ns, not significant; *, $P \leq 0.05$; **, $P \leq 0.001$; ***, $P \leq 0.001$; and ****, $P \leq 0.0001$.

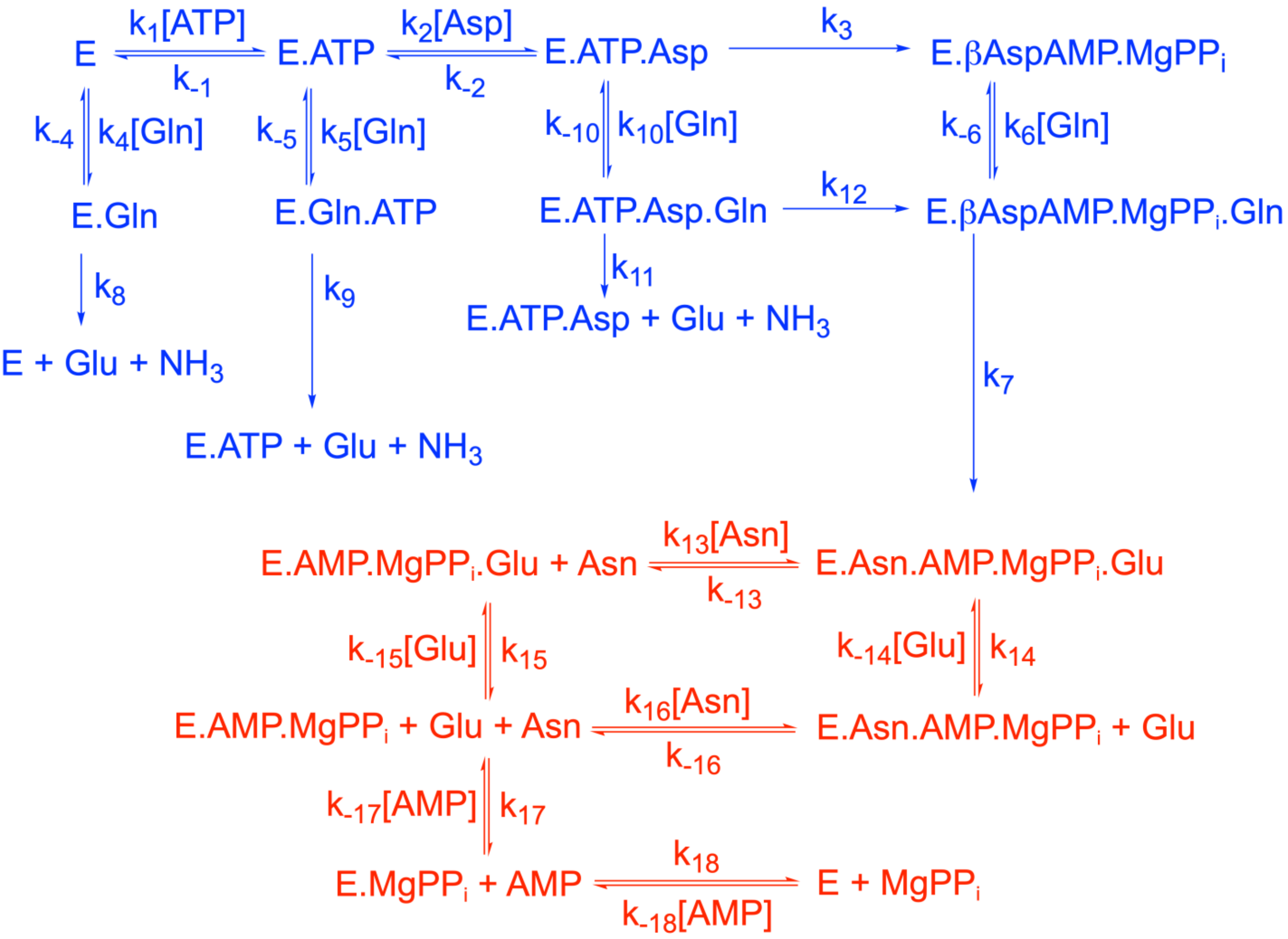


**Supplementary Fig. 6. Putative kinetic mechanism of human ASNS showing intermediate forms of the enzyme that can potentially interact with inhibitors.** This scheme assumes that human ASNS uses the same kinetic mechanism and catalytic intermediates as determined for *Escherichia coli* AS-B. Scheme is based on results reported in (a) Tesson A. R. et al. *Arch. Biochem. Biophys.* **413**, 23-31 (2003) (blue text) and Boehlein, S. K. et al. *Biochemistry* (1999) (red text).

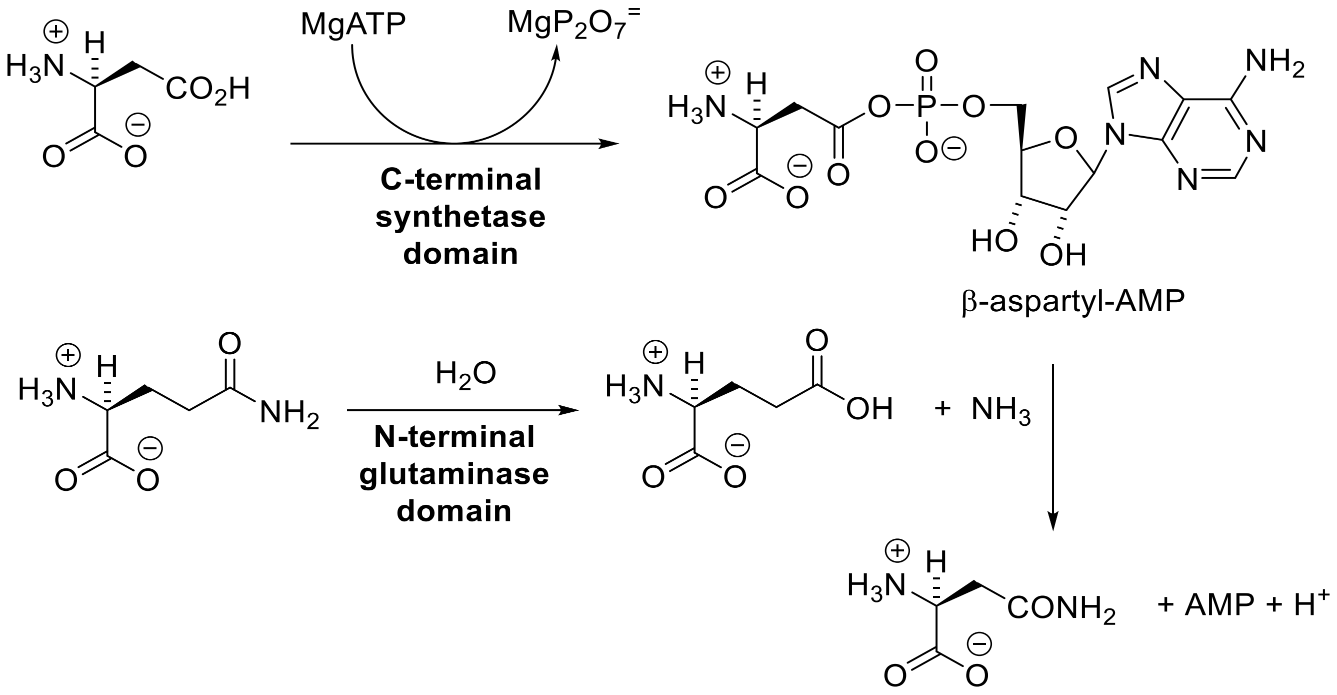


**Supplementary Fig. 7. Reactions catalyzed by the active sites in the N- and C-terminal domains of asparagine synthetase.** Ammonia released in the glutaminase site is translocated to the synthetase site through an intramolecular tunnel linking the two active sites.

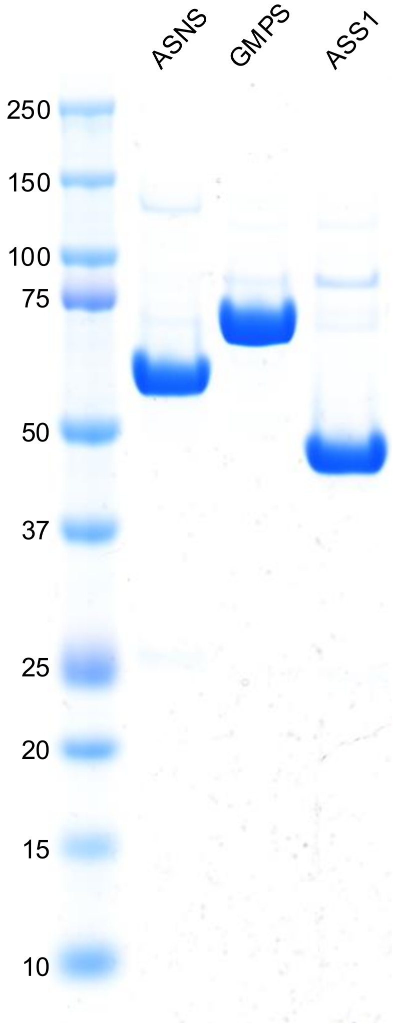


**Supplementary Fig. 8. SDS-PAGE of recombinant human asparagine synthetase (ASNS), GMP synthetase (GMPS) and argininosuccinate synthetase (ASS1) used in this study.** Approximately 4 µg samples of each recombinant protein were subjected to SDS-PAGE and stained with Coomassie Brilliant Blue. Left hand column is composed of molecular weight markers (numbers indicate the marker size in kDa).

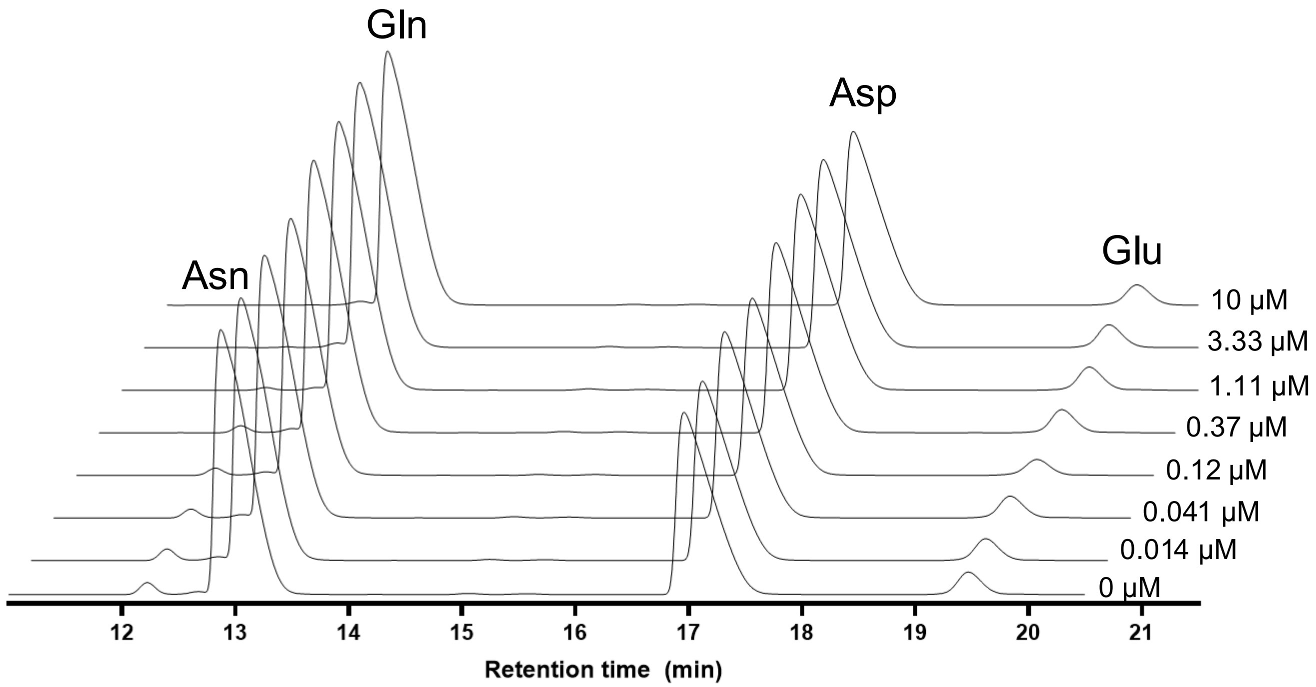


**Supplementary Fig. 9. Representative HPLC traces for inhibition of ASNS-catalyzed asparagine formation by ASX-173.** The effect of titrating the enzyme with ASX-173 in the presence of saturating concentrations of aspartate (Asp), glutamine (Gln) and MgATP is shown by the changes in the area of the asparagine peak (Asn). Concentrations shown on each chromatogram are for ASX-173.

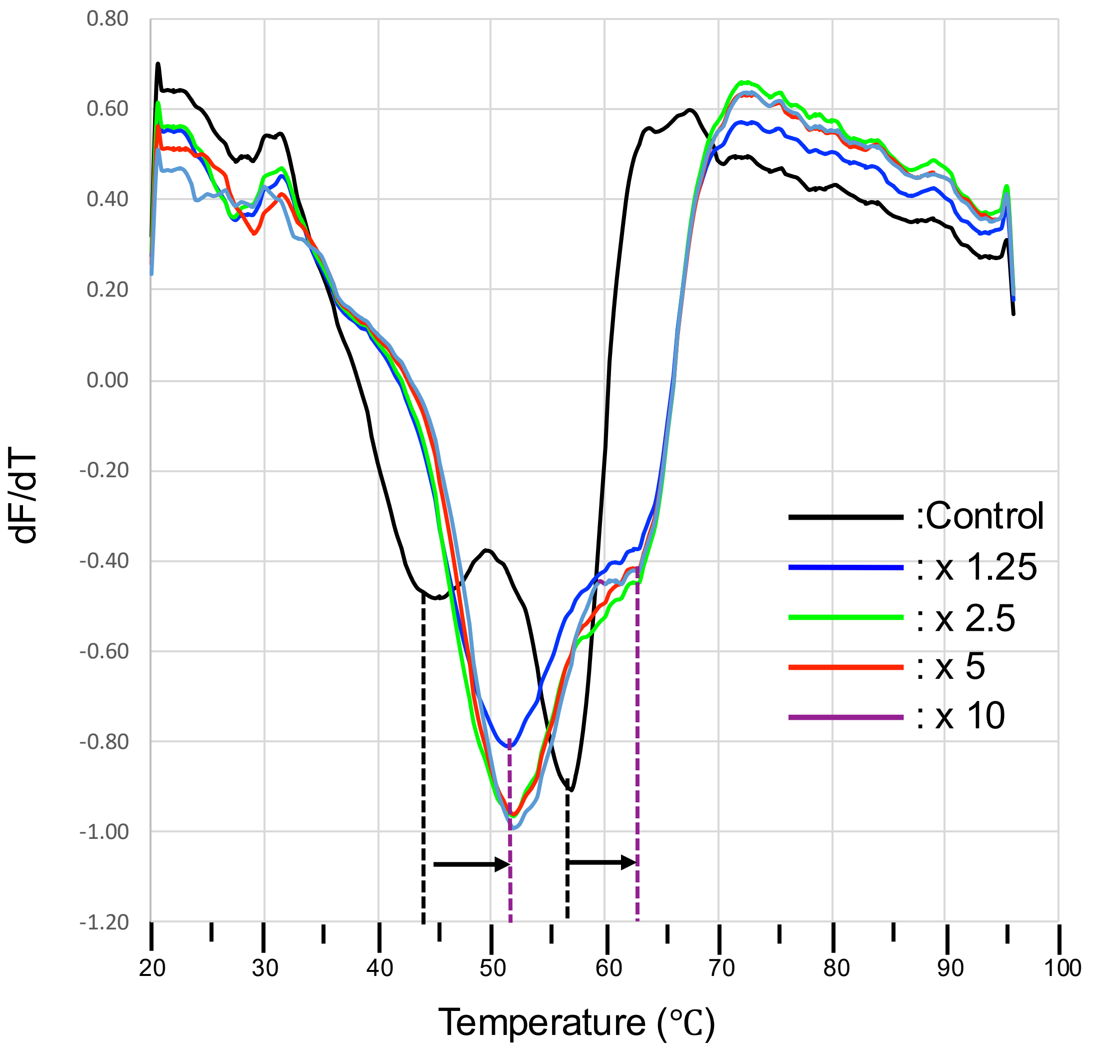


**Supplementary Fig. 10. Titration of ASX-173 for ASNS by thermal shift assay.** The effect of ASX-173 on the thermal stability of recombinant ASNS was assessed by differential scanning fluorimetry (DSF). ASNS (6 µg per assay) was incubated at room temperature for 1 h with varying molar ratios of ASX-173 (0 [control], 1.25, 2.5, 5.0, and 10) in the presence of 5 mM ATP and 10 mM $MgCl_2$ prior to DSF measurement. First-derivative melting curves are shown for each condition: control (black), 1.25× (blue), 2.5× (green), 5.0× (red), and 10× (purple). A dotted line highlights the Tm for control (black) and ASX-173 at all molar ratios (purple), with the Tm shift indicated by arrows.

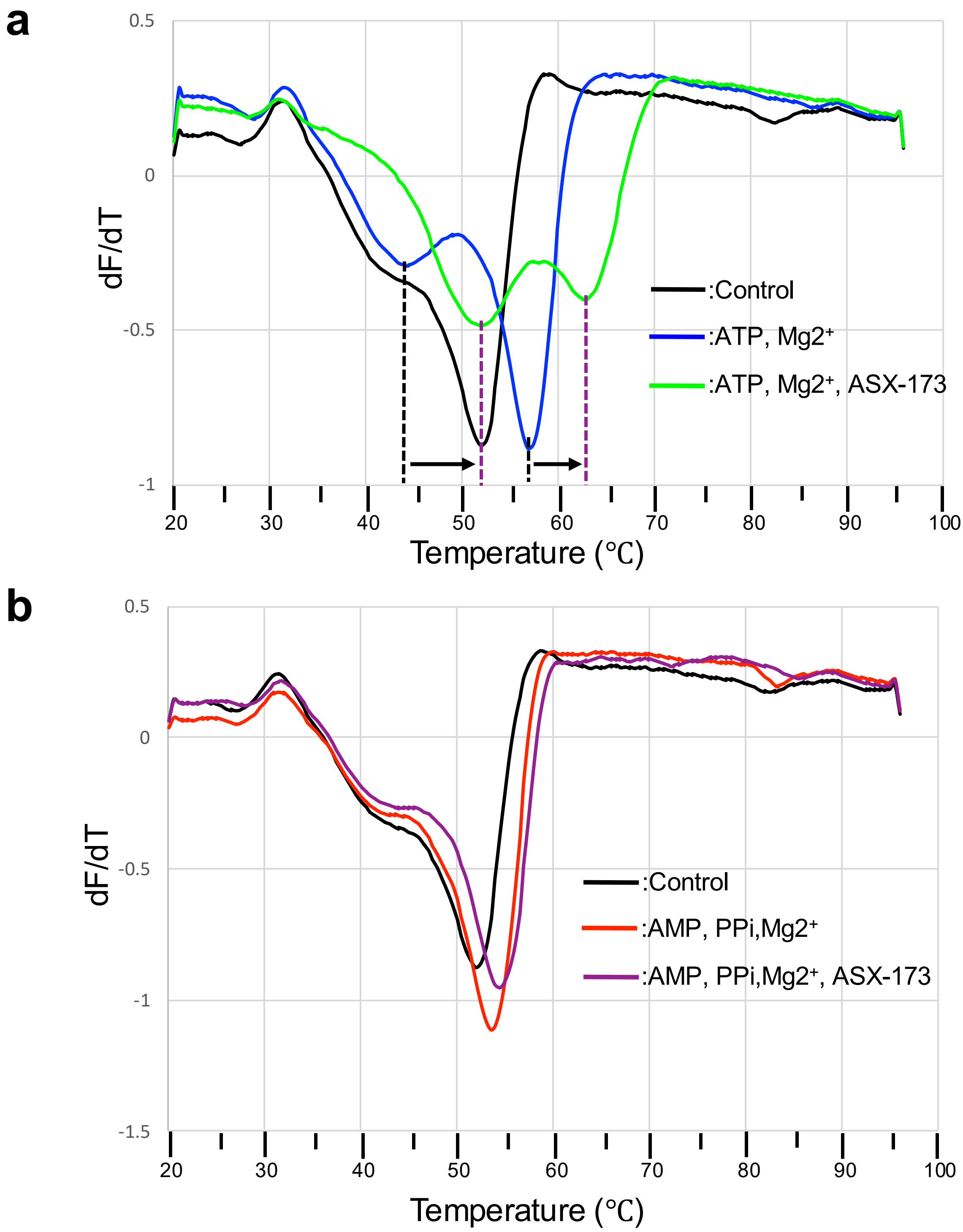


**Supplementary Fig. 11. Effect of ASX-173 on the thermal stability of ASNS in the presence of AMP, PPi, and $Mg^{2+}$.** The impact of ASX-173 on the thermal stability of recombinant ASNS was evaluated by differential scanning fluorimetry (DSF). ASNS (6 µg per assay) was incubated at room temperature for 1 h under six different conditions: (a) Control (buffer only, black), 5 mM ATP with 10 mM $Mg^{2+}$ (blue), and 5 mM ATP with 10 mM $Mg^{2+}$ plus ASX-173 at 10-fold molar excess (green). A dotted line highlights the Tm for the control (black) and ATP/$Mg^{2+}$ plus ASX-173 condition (green), with the Tm shift indicated by arrows. (b) Control (buffer only, black), 5 mM AMP with 5 mM PPi and 10 mM $Mg^{2+}$ (red), and 5 mM AMP with 5 mM PPi and 10 mM $Mg^{2+}$ plus ASX-173 at 10-fold molar excess (purple).

**Table 1. Cryo-EM data collection, refinement and validation statistics**

| **Data collection** | | |
|---|---|---|
| | ASNS-ASX173 | ASNS-ASX173 |
| Magnification | ×150,000 | ×150,000 |
| Voltage (kV) | 200 | 300 |
| Electron exposure (e/Å$^2$) | 40 | 59.45 |
| Defocus range (μm) | -0.4 to -2.0 | -0.8 to -1.8 |
| Pixel size (Å) | 0.93 | 0.411 |
| **Data processing** | | |
| | **EMD-73230, PDB 9YNS** | **EMD-73232, PDB 9YNT** |
| Symmetry imposed | C2 | C2 |
| Initial particle images (no.) | 4,151,760 | 5,813,013 |
| Final particle images (no.) | 88,344 | 117,146 |
| Map resolution (Å) | 3.14 | 2.58 |
| FSC threshold | 0.143 | 0.143 |
| Map resolution range | 2.4-3.6 | 2.0-3.2 |
| Map sharpening B factor (Å$^2$) | 104.2 | 98.2 |
| **Refinement** | | |
| Model resolution (Å$^2$) | 2.18 | 1.97 |
| Model composition | | |
| Non-hydrogen atoms | 8379 | 8289 |
| Protein residues | 1035 | 1027 |
| Ligands: | | |
| Mg | 4 | 4 |
| AMP | 2 | 2 |
| PPi | 2 | 2 |
| ASX-173 | 2 | 2 |
| B-factors (min/max/mean) | | |
| Protein | 22.01/112.93/48.85 | 17.45/108.41/41.80 |
| Ligand | 6.45/75.39/51/51 | 25.25/48.17/35.95 |
| r.m.s.d. deviations | | |
| Bond length (Å) | 0.003 | 0.003 |
| Bond angles (° ) | 0.543 | 0.524 |
| Validation | | |
| MolProbility score | 2.36 | 2.27 |
| Clashscore | 10.09 | 9.02 |
| Rotamer outliers (%) | 4.28 | 4.52 |
| CaBLAM outliers (%) | 3.2 | 2.7 |
| Ramachandran plot (%) | | |
| Outliers | 0 | 0 |
| Allowed | 5.11 | 4.23 |
| Favored | 94.89 | 95.77 |

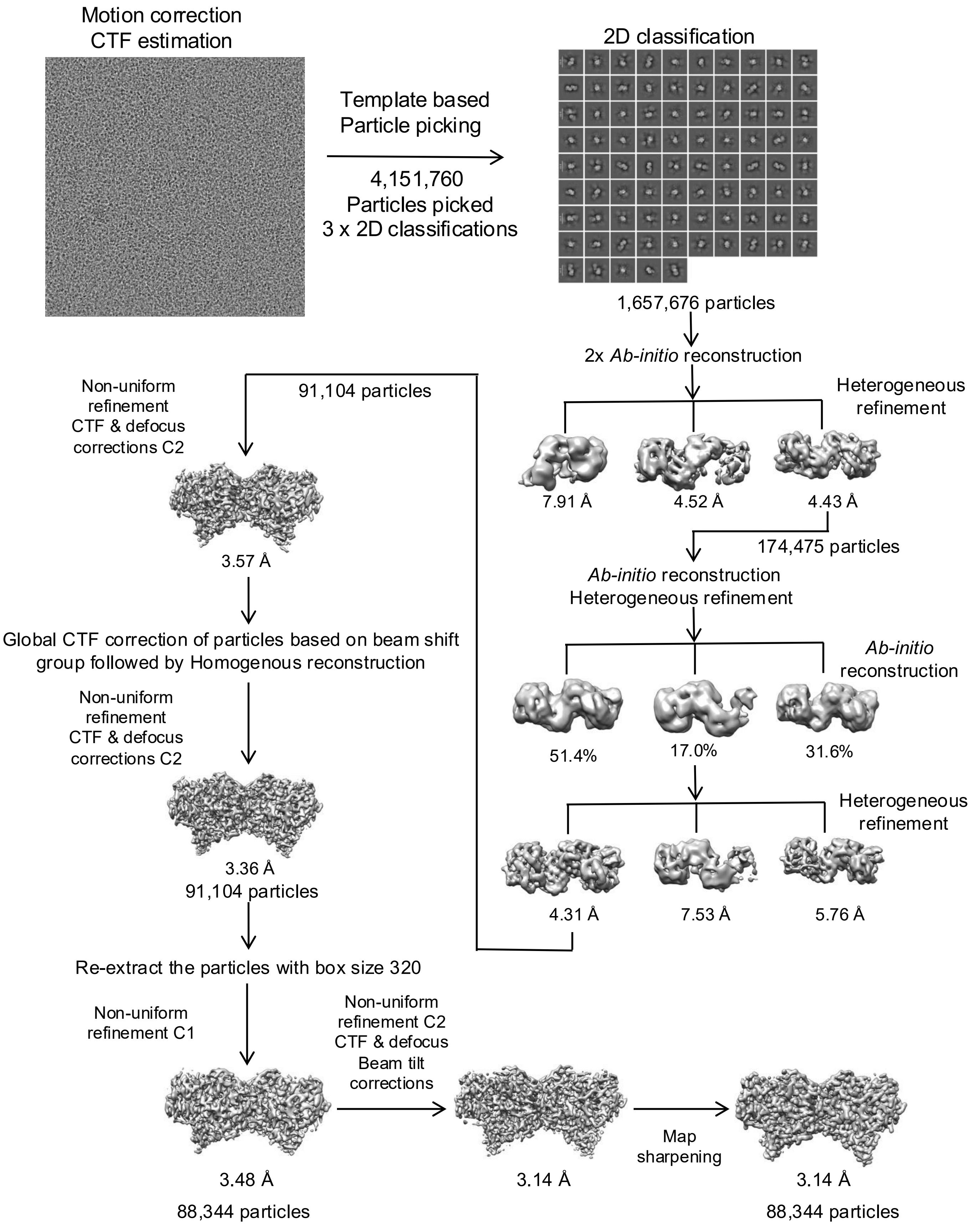


**Supplementary Fig. 12. Summary of the cryo-EM data processing workflow of 200 kV cryo-EM dataset using cryoSPARC v4.7.0.**

**a Gold standard FSC curve for ASNS-ASX173 complex by 200kV cryo-EM**

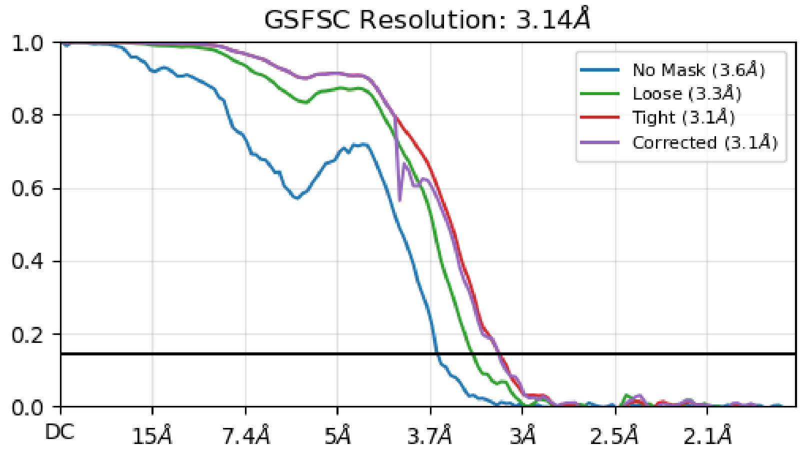


**b Viewing distribution plots**

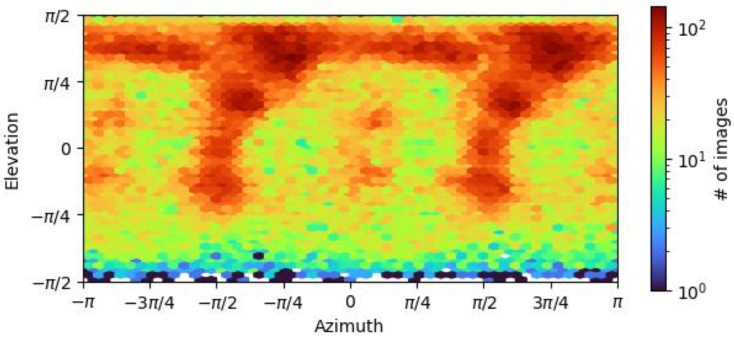


**c cFSC and cFAR**

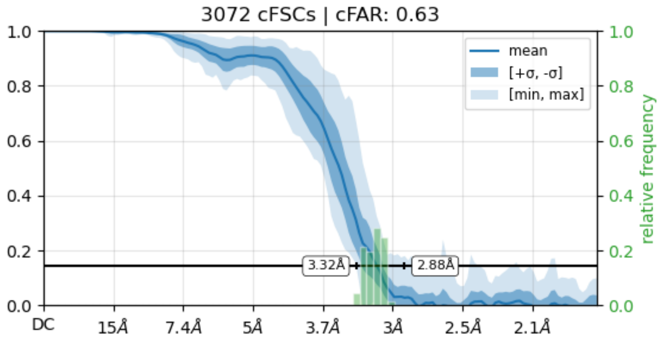


**d Local resolution**

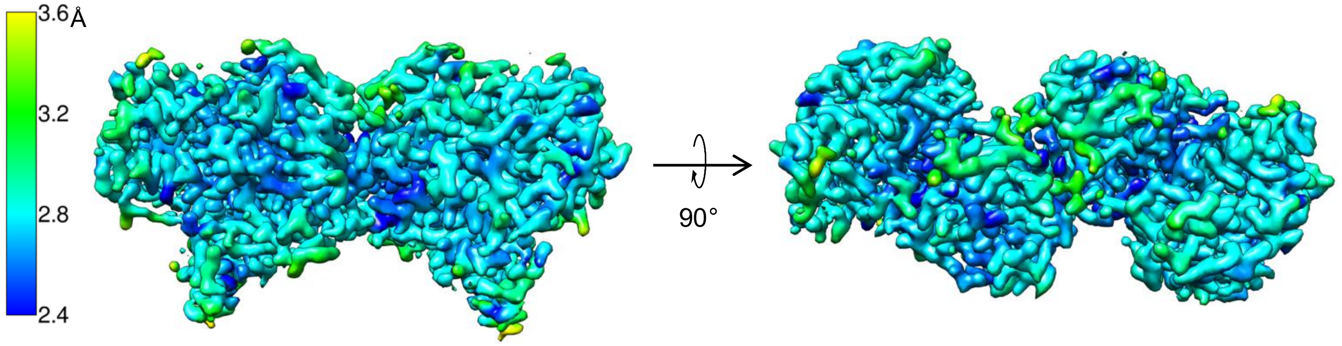


**Supplementary Fig. 13. Assessment of the quality of cryo-EM map determined by 200kV scope.**
**a** Gold standard FSC curve for the map of ASNS-ASX173 complex generated in cryoSPARC v4.7.0
**b** Viewing distribution plots for the map generated in cryoSPARC v4.7.0
**c** Measure of anisotropy is indicated by cFSC and cFAR
**d** Local resolution of the map (color scale shown on the left).

### a Representative densities and corresponding models for WT human ASNS

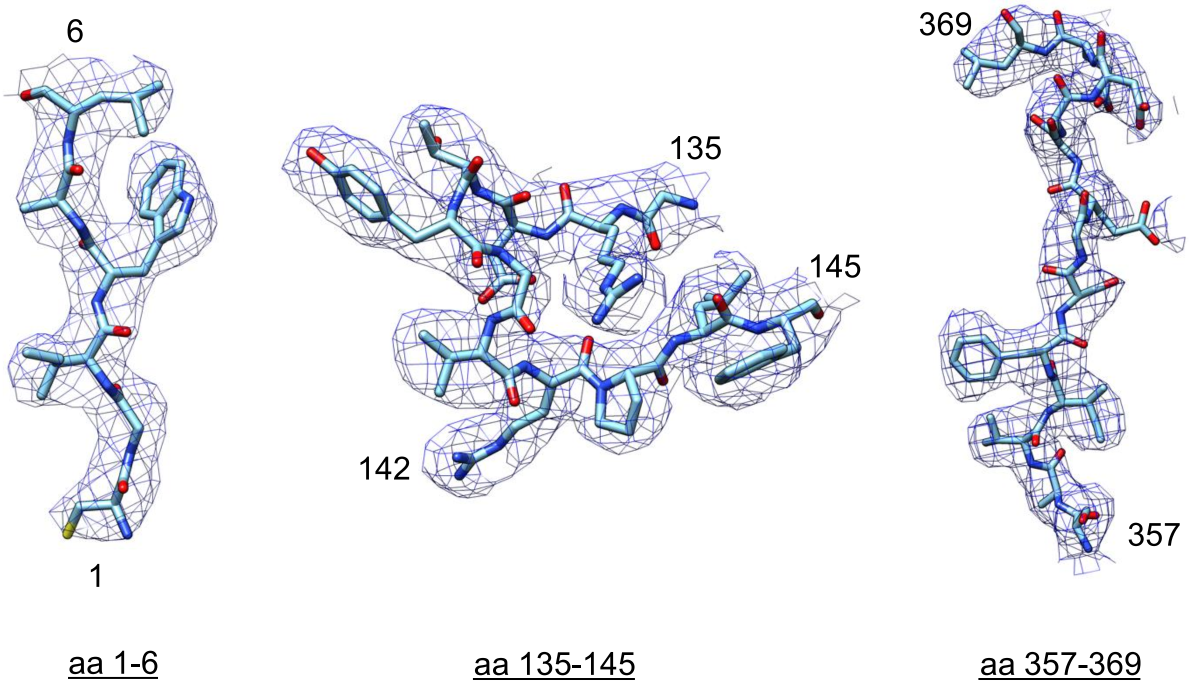


### b Map-model FSC curve

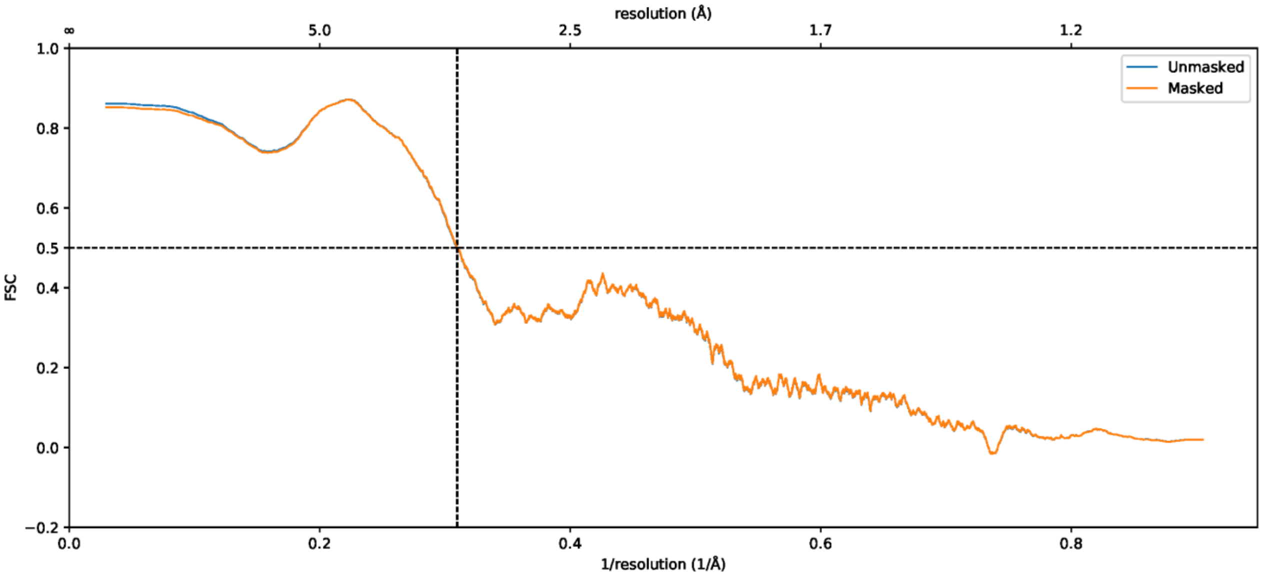


**Supplementary Fig. 14. Validating the cryo-EM structure: map and its model by 200 kV cryo-EM**
**a** Representative density of the map for residues in the N-terminal catalytic region (aa 1-6), the ammonia tunnel region (aa 132-145), and the C-terminal active site (aa 357-369)
**b** Model-to-map FSC curves generated in Phenix.

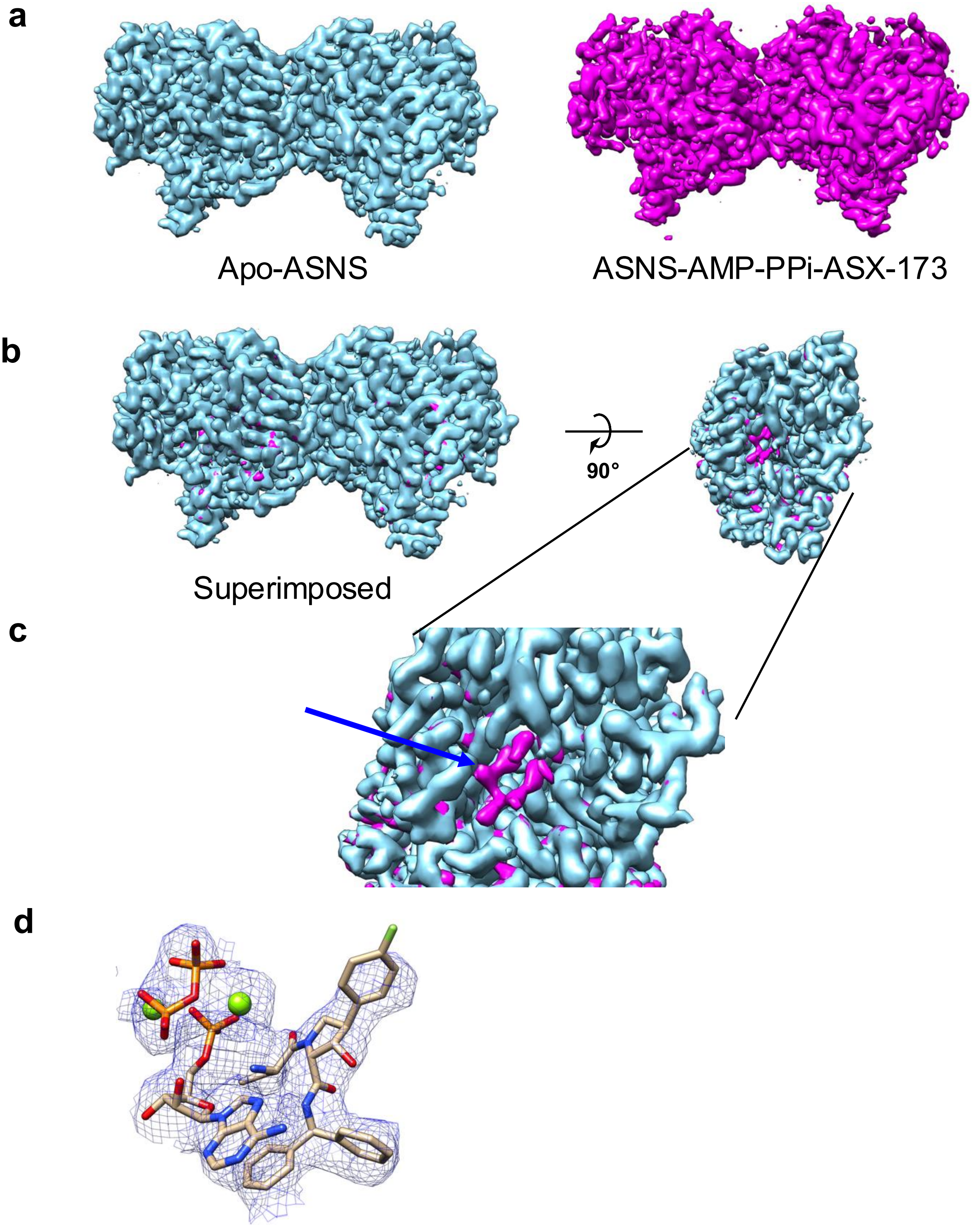


**Supplementary Fig. 15. Extra density observed in the ASNS–ASX-173 complex in the presence of ATP and $Mg^{2+}$ a** Cryo-EM map of apo-ASNS (EMD-40764, sky blue) (left) compared with the map of the ASNS–ASX-173 complex in the presence of ATP and $Mg^{2+}$ (magenta, right). **b** Superimposed maps of apo- and ASX-173–bound ASNS in front view (left) and side view (right). **c** Close-up view of the C-terminal synthetase active site highlighting the extra density observed in the ASX-173–bound state (blue arrow). **d** Docking of ASX-173, AMP, pyrophosphate (PPi), and $Mg^{2+}$ ions into the extra density identified from the ASX-173–bound map, generated from datasets collected on a 200 kV Glacios microscope. $Mg^{2+}$ colored in green, PPi (red, orange)

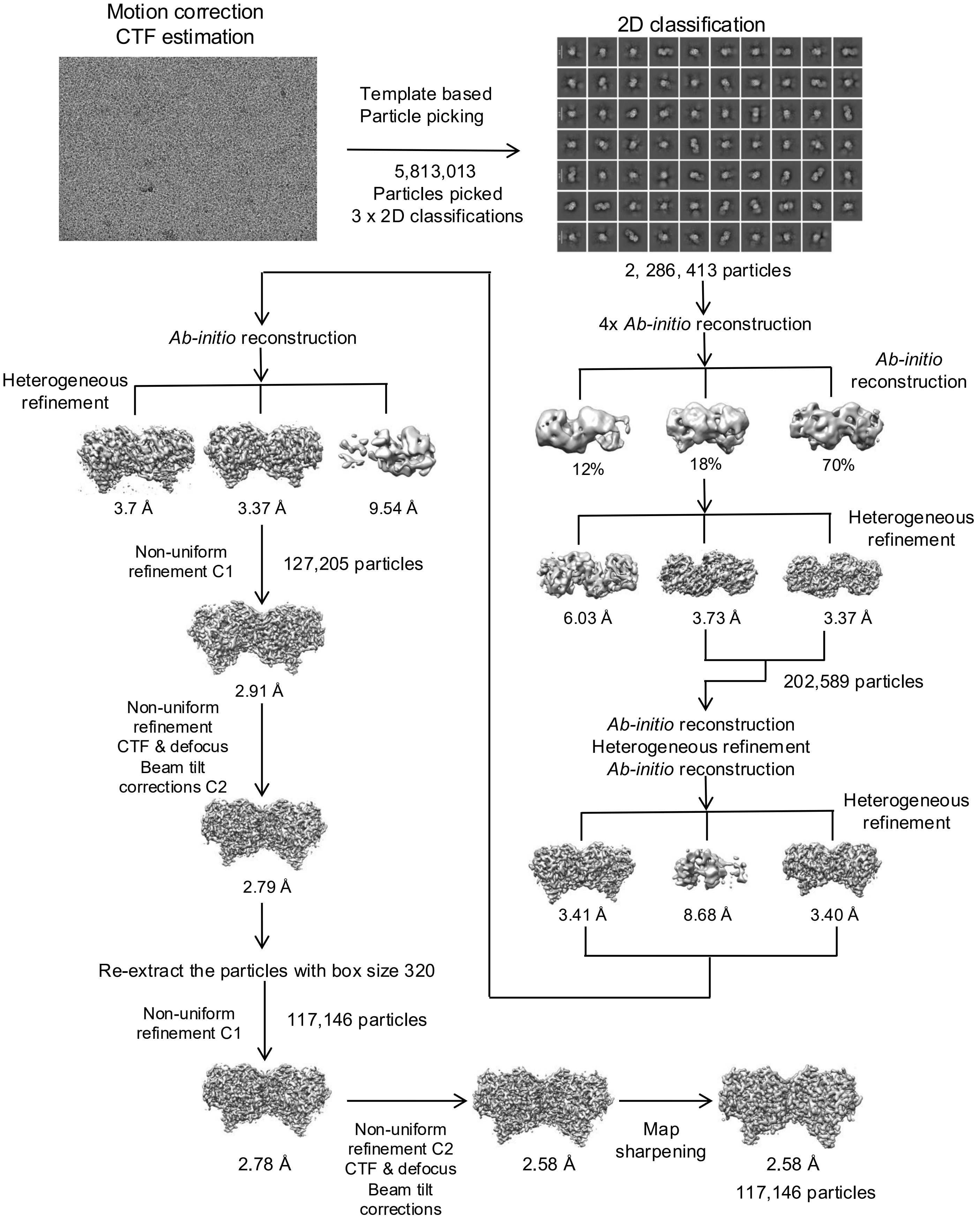


**Supplementary Fig. 16. Summary of the cryo-EM data processing workflow of 300 kV cryo-EM dataset using cryoSPARC v4.7.0.**

**a Gold standard FSC curve for ASNS-ASX173 complex by 300 kV cryo-EM**

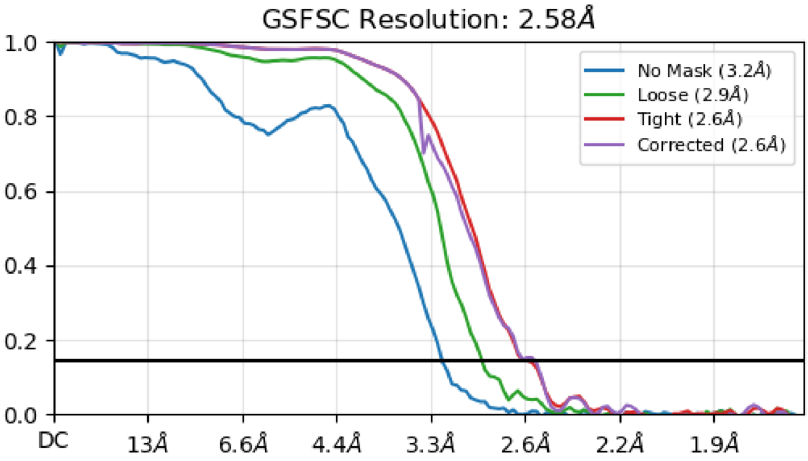


**b Viewing distribution plots**

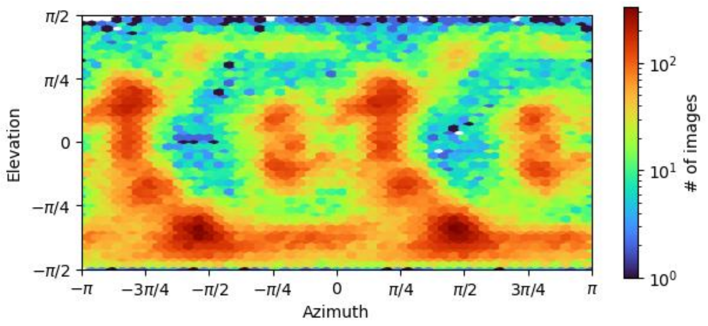


**c cFSC and cFAR**

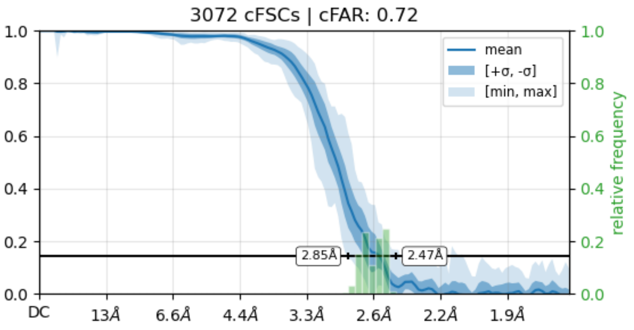


**d Local resolution**

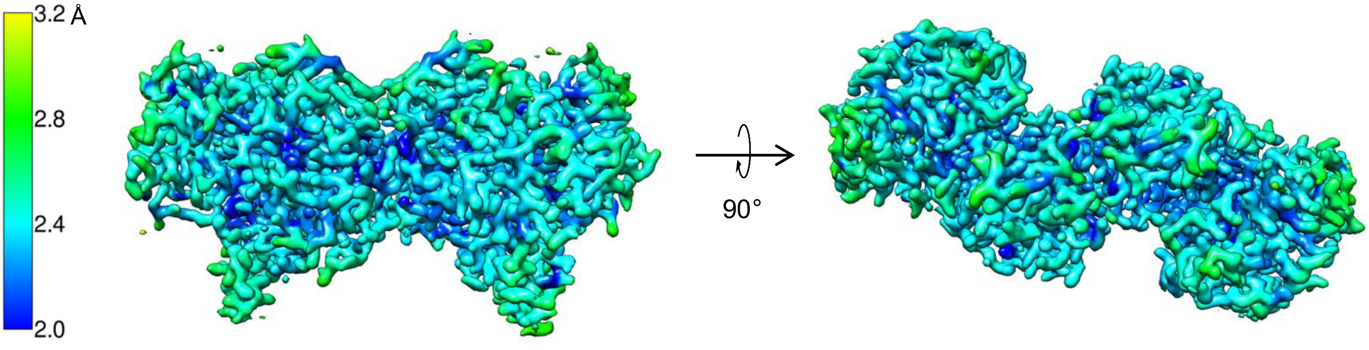


**Supplementary Fig. 17. Assessment of the quality of cryo-EM map determined by 300 kV cryo-EM**

**a** Gold standard FSC curve for the map of ASNS-ASX173 complex generated in cryoSPARC v4.7.0

**b** Viewing distribution plots for the map generated in cryoSPARC v4.7.0

**c** Measure of anisotropy is indicated by cFSC and cFAR

**d** Local resolution of the map (color scale shown on the left).

### a Representative densities and corresponding models for ASNA-ASX173

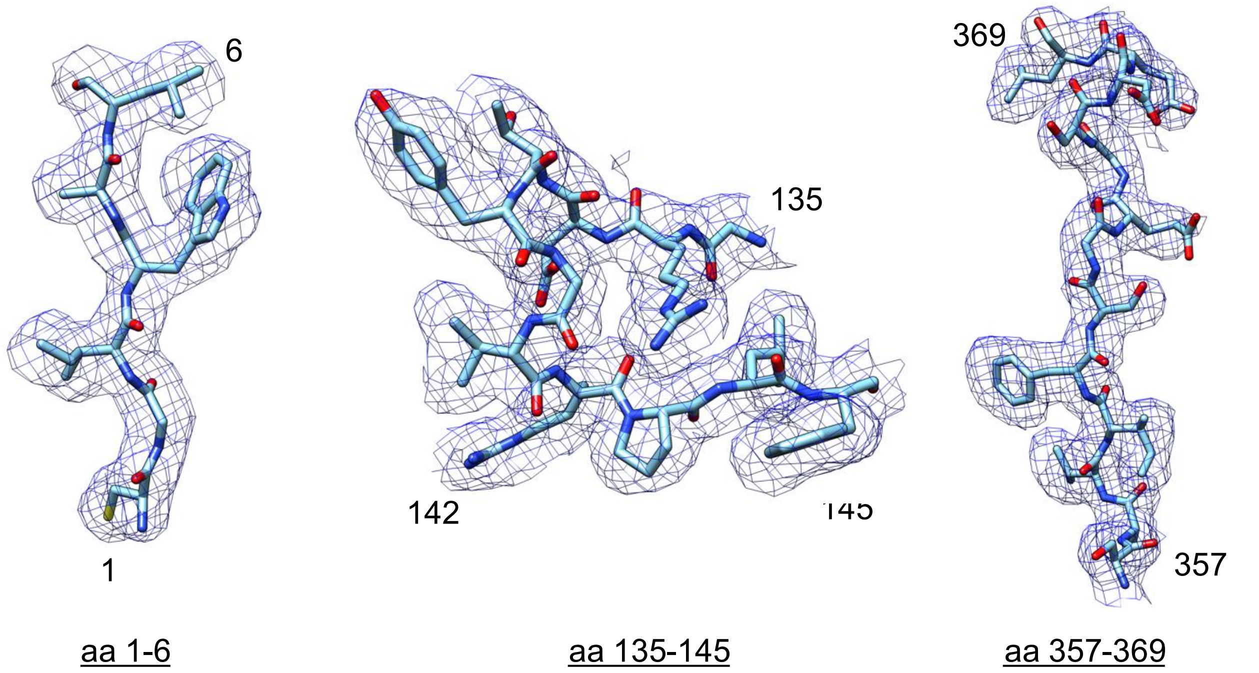


### b Map-model FSC curve

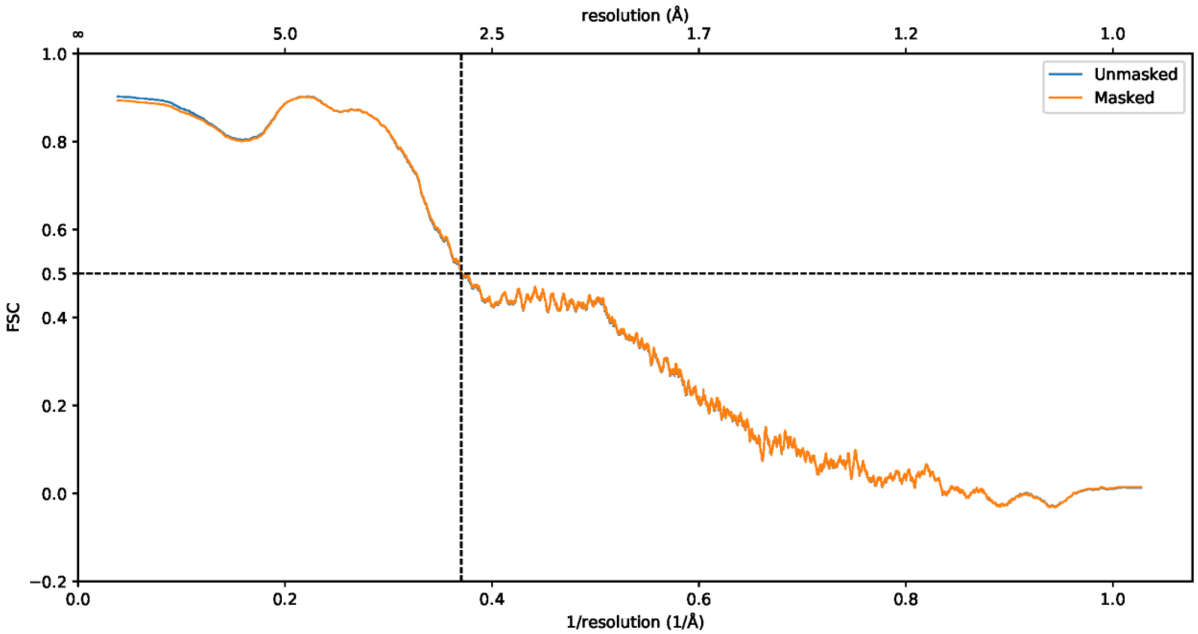


**Supplementary Fig. 18. Validating the cryo-EM structure: map and its model by 300 kV cryo-EM**

**a** Representative density of the map for residues in the N-terminal catalytic region (aa 1-6), the ammonia tunnel region (aa 132-145), and the C-terminal active site (aa 357-369)

**b** Model-to-map FSC curves generated in Phenix.

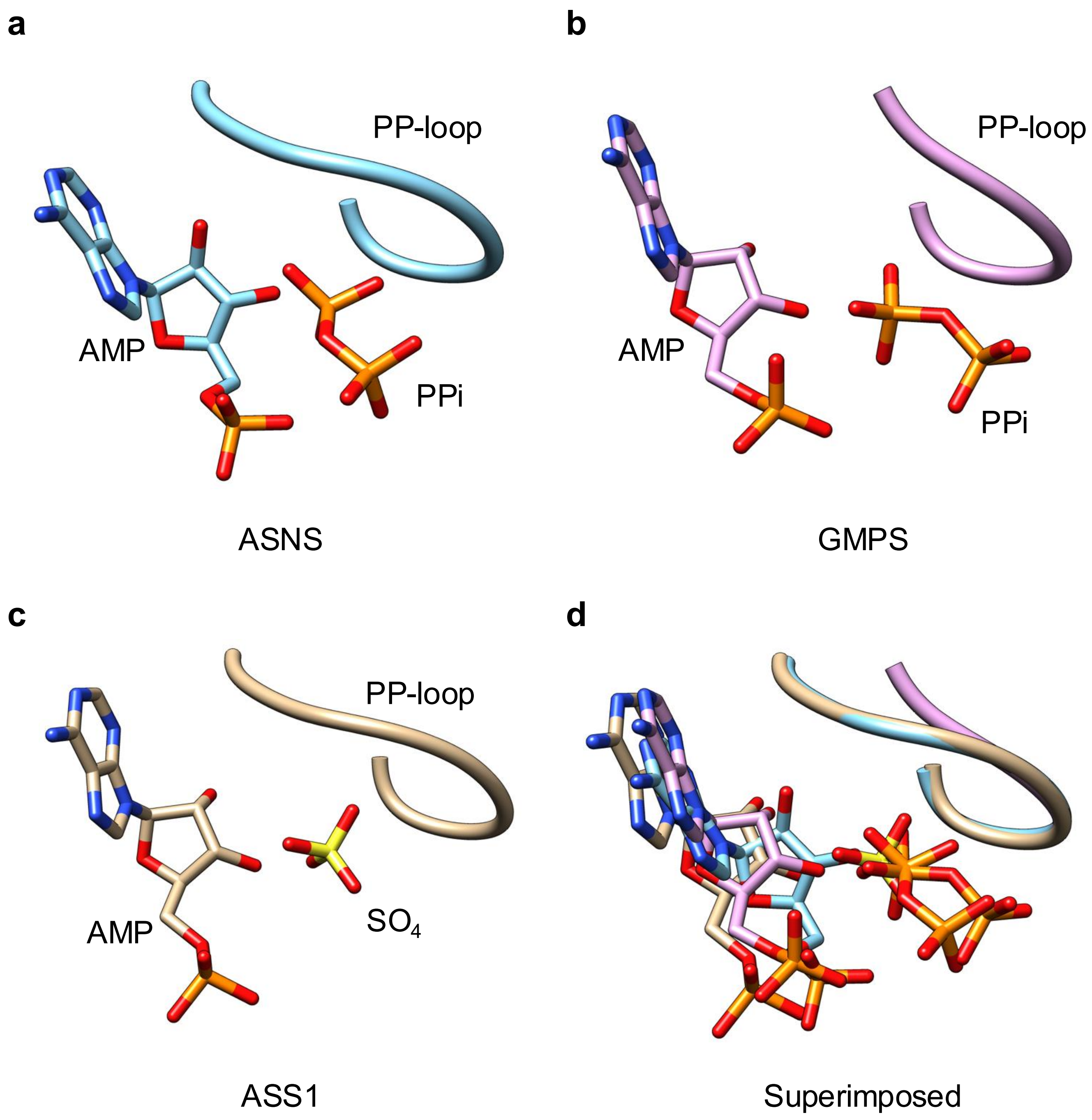


**Supplementary Fig. 19. AMP–P-loop comparison among ASNS, GMPS, and ASS1.** **a** ASNS–AMP–PPi–ASX-173 complex (PDB: xx), **b** human GMPS (PDB: 1GPM), and **c** *T. thermophilus* ASS1. P-loops are shown as ribbons (ASNS, sky blue; GMPS, pink; ASS1, goldenrod). Sulfate ions are in yellow. d Superimposition highlights structural similarity (RMSD: 0.155 Å, ASNS vs. GMPS; 0.240 Å, ASNS vs. ASS1).

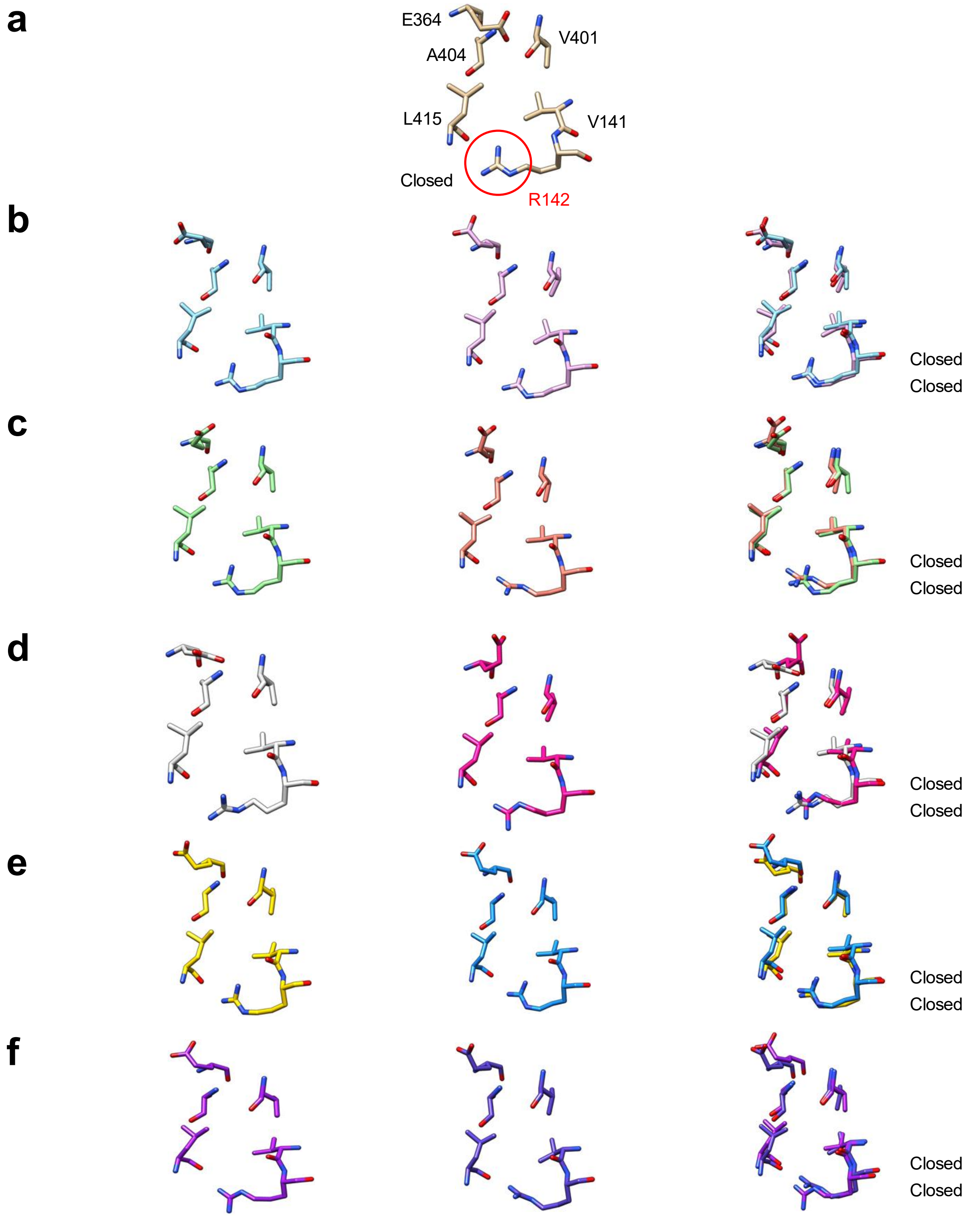


**Supplementary Fig. 20. PCA-derived conformations of the ammonia tunnel in the ASNS–AMP–PPi–ASX-173 complex obtained from 3D variability analysis (3DVA).** (a) Side view of key residues (Val141, Arg142, Glu364, Val401, Ala404, Val414, Leu415) forming the ammonia tunnel in the EM map of apo-ASNS (EMD-40764). The side chain of Arg142, which blocks the tunnel, is highlighted (red circle). (b–f) Side views of the same residues in variable EM maps from 3DVA of the ASNS–AMP–PPi–ASX-173 complex, with corresponding models generated by 3D variability refinement. Residues from the tunnel conformations at frame 0 (one end) and frame 19 (the opposite end) are superimposed for each PCA component: (b) component 0, (c) component 1, (d) component 2, (e) component 3, (f) component 4.

**a**

**b**

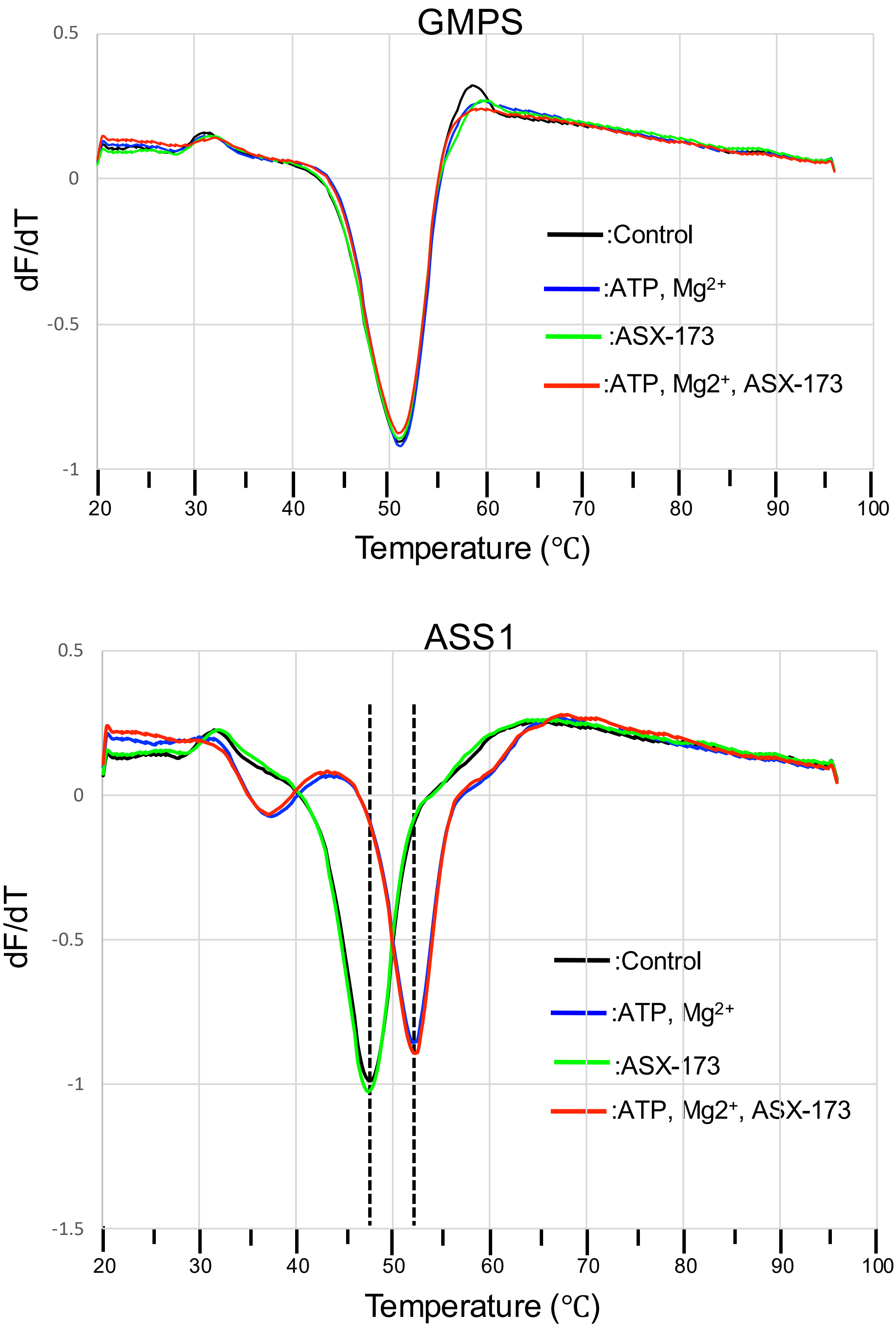


**Supplementary Fig. 21. Effect of ASX-173 on human GMP synthetase (GMPS) and arginine-succinate synthetase (ASS1) by thermal shift assays.** Thermal stability of recombinant (a) human GMPS (6 µg per assay) and (b) human ASS1 (8 µg per assay) was analyzed by differential scanning fluorimetry. First-derivative melting curves are shown, with melting temperatures (Tm) indicated for each condition: control (buffer only, black), ATP and $Mg^{2+}$ (blue), ASX-173 at 10-fold molar excess (green), and ATP, $Mg^{2+}$, plus ASX-173 at 10-fold molar excess (red).

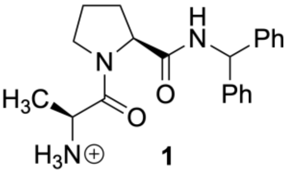

**1**

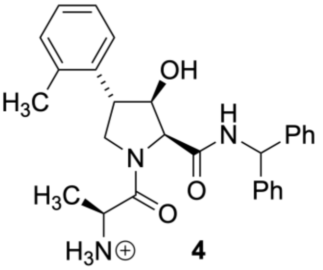

**4**

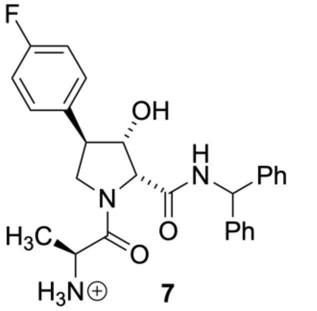

**7**

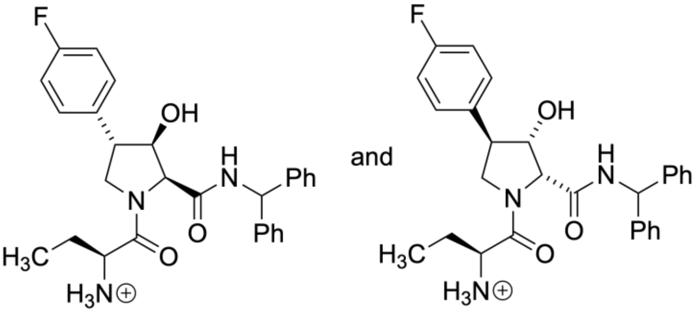

**15** (mixture of diastereoisomers)

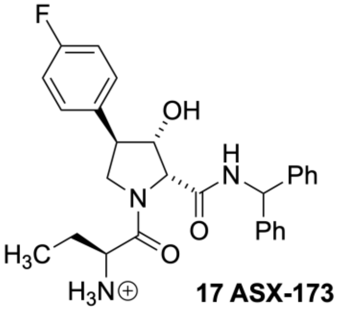

**17 ASX-173**

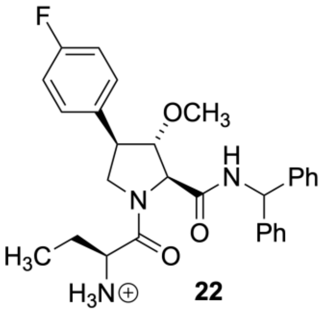

**22**

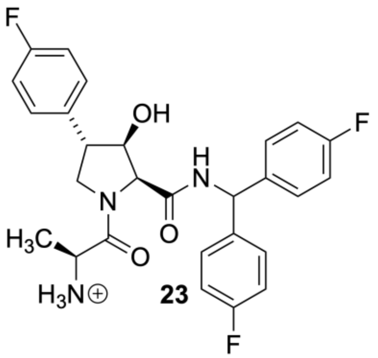

**23**

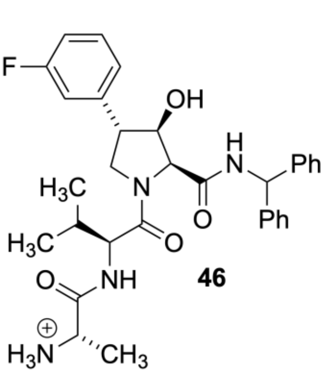

**46**

**Supplementary Fig. 22**. **Chemical structures of ASX-173 and selected analogs**. Compound numbers correspond to those given in a patent (Genis et al., *Compounds that inhibit asparagine synthetase and methods of use*: WO 2021/236475A1).

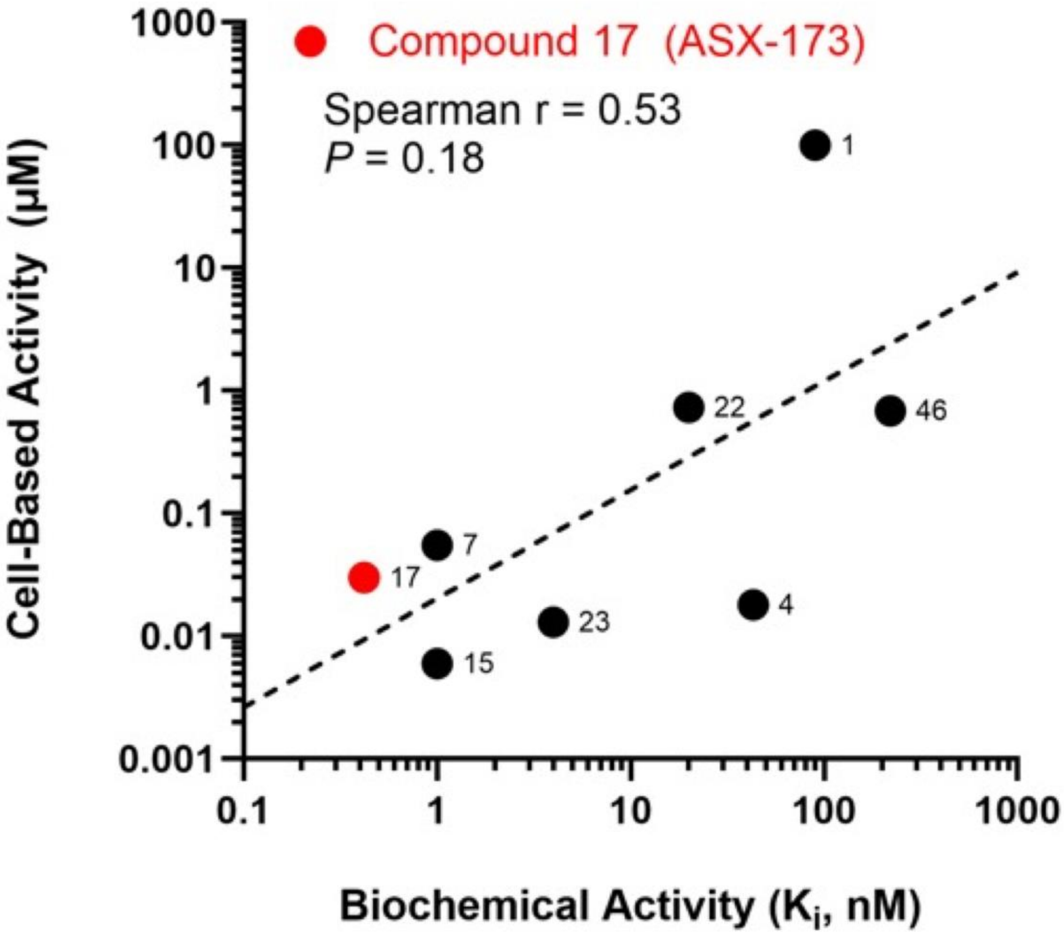


**Supplementary Fig. 23. Correlation between the ability of ASX-173 and selected analogs to inhibit the ammonia-dependent activity of human ASNS ($K_i$*) and cytotoxicity ($IC_{50}$) in a Jurkat T cell proliferation assay.** Data are taken from a patent (Genis et al., *Compounds that inhibit asparagine synthetase and methods of use*: WO 2021/236475A1). Compound numbers correspond to those given in the patent and are also shown in Supplementary Fig. S19. Values for compound 17 (ASX-173) are indicated in red.

### Table 2. Primers used for qRT-PCR analysis

| Gene | Forward Primer | Reverse Primer |
|---|---|---|
| ATF4 | 5'-TCAAACCTCATGGGTTCTCC-3' | 5'-GTGTCATCCAACGTGGTCAG-3' |
| ASNS | 5'-TAACGCGTGCGGGAAGTT-3' | 5'-CCTGTGCGCGCTGGTT-3' |
| CHOP | 5'-AGCCAAAATCAGAGCTGGAA-3' | 5'-ACAAGTTGGCAAGCTGGTCT-3' |
| GADD34 | 5'-TCCGACTGCAAAGGCGGCTCA-3' | 5'-CAGCCAGGAAATGGACAGTGAC-3' |
| TRIB3 | 5'-GCTTTGTCTTCGCTGACCGTGA-3' | 5'-CTGAGTATCTCAGGTCCCACGT-3' |
| Actin | 5'-GGACTTCGAGCAAGAGATGG-3' | 5'-AGCACTGTGTTGGCGTACAG-3' |
